# The influence of feeding behaviour and temperature on the capacity of mosquitoes to transmit malaria

**DOI:** 10.1101/604249

**Authors:** Eunho Suh, Marissa K. Grossman, Jessica L. Waite, Nina L. Dennington, Ellie Sherrard-Smith, Thomas S. Churcher, Matthew B. Thomas

**Affiliations:** Center for Infectious Disease Dynamics, Department of Entomology, Penn State University, University Park, PA 16802; Green Mountain Antibodies, PO Box 1283, Burlington, VT 05402; MRC Centre for Global Infectious Disease Analysis, Department of Infectious Disease Epidemiology, Imperial College London, Norfolk Place, London W2 1PG, UK

**Author notes:** Correspondence to be sent to: Eunho Suh, Center for Infectious Disease Dynamics, Department of Entomology, Penn State University, University Park, PA 16802.

**Keywords:** Thermal ecology, diurnal temperature variability, *Anopheles* mosquitoes, *Plasmodium* malaria, biting behaviour, behavioral resistance, residual transmission

## Abstract

Insecticide-treated bed nets reduce malaria transmission by limiting contact between mosquito vectors and human hosts when mosquitoes feed during the night. However, malaria vectors can also feed in the early evening and in the morning when people are not protected. Here, we explored how timing of blood feeding interacts with environmental temperature to influence the capacity of *Anopheles* mosquitoes to transmit the human malaria parasite, *Plasmodium falciparum*. We found no effect of biting time itself on the proportion of mosquitoes that became infectious (vector competence) at constant temperature. However, when mosquitoes were maintained under more realistic fluctuating temperatures there was a significant increase in competence for mosquitoes feeding in the evening, and a significant reduction in competence for those feeding in the morning, relative to those feeding at midnight. These effects appear to be due to thermal sensitivity of malaria parasites during the initial stages of parasite development within the mosquito, and the fact that mosquitoes feeding in the evening experience cooling temperatures during the night, whereas mosquitoes feeding in the morning quickly experience warming temperatures that are inhibitory to parasite establishment. A transmission dynamics model illustrates that such differences in competence could have important implications for disease endemicity, the extent of transmission that persists in the presence of bed nets, and the epidemiological impact of behavioural resistance. These results indicate the interaction of temperature and feeding behaviour to be a major ecological determinant of the vectorial capacity of malaria mosquitoes.

## Introduction

Wide-scale use of long-lasting insecticide-treated bed nets (LLINs) and indoor residual insecticide sprays (IRS) has led to substantial declines in the global burden of malaria in recent years^1^. However, these gains are now threatened by the evolution of insecticide resistance^2–4^. Studies from many locations demonstrate both target site and metabolic resistance to be widespread in malaria vector populations^2–4^. In addition, there are growing reports of behavioural resistance, such as changes in mosquito biting behaviour (i.e. “anti-insecticide” behaviour), which reduce the probability of insecticide encounter and/or attenuate the efficacy of insecticides^5–8^.

In principle, physiological mechanisms of resistance can be countered by switching classes of insecticide, or using synergists to disrupt detoxification mechanisms^9–13^. However, behavioural resistance is potentially more insidious since changes in biting time (e.g. early evening biting before humans are protected under bed nets) and/or shifts in biting location (outdoor biting rather than indoors) could render whole classes of vector control tools ineffective^5–8, 14^. Furthermore, even in the absence of behavioural or physiological resistance, typical biting patterns for many malaria vectors still span periods of the evening and morning, when effective coverage of bed nets is less^15, 16^. This crepuscular biting behaviour contributes to ‘residual transmission,’ which is defined as the transmission that persists after achieving full universal coverage with an effective intervention such as LLINs, to which local vector populations are fully susceptible ^15–17^.

Vector competence describes the ability of an arthropod to become infected, allow replication, and ultimately transmit a pathogen^18^. In order to become transmissible, malaria parasites go through multiple developmental stages within the mosquito, progressing from the gametocytes ingested in the blood meal, to gametes, the fertilized zygotes, the motile ookinetes that invade the mosquito midgut, the oocyst in which the parasite undergoes replication, and finally to the sporozoites that invade the salivary glands and can be passed onto a new host during a subsequent blood meal^19, 20^. Competence is determined by both genetic and environmental factors^18, 21^. Mosquito gene expression is known to follow circadian rhythms^22–24^. Further, temperatures in many malaria endemic areas exceed 30°C as temperatures fluctuate during the day ^25–28^, and early parasite infection is known to be sensitive to high temperatures^29, 30^. These extrinsic and intrinsic factors could have direct or indirect effects on parasite survival and establishment and hence, contribute to variation in competence of mosquitoes feeding at different times of the day^23, 29–31^. Understanding any such variation is key to fully understanding transmission ecology.

Here, we explore the effect of time-of-day of feeding on vector competence of *Anopheles* mosquitoes to determine whether all mosquitoes are equally capable of transmitting malaria, and to better understand the potential epidemiological consequences of shifts in feeding behaviour. First, we review recent literature to characterise biting activity of *Anopheles* mosquitoes in the field. We find many examples indicating peak biting to occur in the evening or morning rather than the middle of the night, as well as evidence to suggest recent changes in biting time following wide-scale distribution of bed nets. We next use a series of laboratory infection studies to examine whether timing of blood feeding affects vector competence, considering both intrinsic (circadian) and extrinsic (temperature) factors. We find that while there is little apparent effect of circadian rhythm alone, diurnal temperature fluctuation leads to a significant increase in the vector competence of evening biting mosquitoes, but a decrease for morning biting mosquitoes. To explore the possible epidemiological implications of this variation in competence we use a mathematical model of malaria transmission. This model analysis suggests that differences in vector competence associated with the interaction of temperature and mosquito biting behaviour could have a noticeable impact on disease endemicity, and alter the relative efficacy of LLINs. Finally, we conduct a further set of experiments to begin to elaborate on the mechanisms underpinning the variation in competence and to determine whether the effects might be mitigated by mosquito thermal behaviour. These experiments suggest that the changes in vector competence are associated with high thermal sensitivity of the parasites during the initial infection process, and are likely robust as mosquitoes appear behaviourally unresponsive to temperatures that are critically damaging to parasite establishment. Overall, our results suggest the interaction of biting time and temperature to be a major ecological driver of vectorial capacity.

## Results

### Daily biting activity of malaria mosquitoes

We reviewed the contemporary malaria control literature published between 2000 and 2017 using PubMed to examine the biting activity of *Anopheles* mosquitoes. The goal of this study was to characterise biting activity of mosquitoes during the period in which the use of LLINs in sub-Saharan Africa was scaled up substantially^1^. We identified 270 papers that referred to biting time of malaria vectors, with 42 papers providing measures of hourly biting activity. Peak biting time of most malaria vectors is generally considered to occur around 00:00-04:00h^15, 32^ and from these 42 papers, we identified 78 cases where biting conformed to this conventional pattern (Supplementary Table 1 and Supplementary Table 2). However, we identified 64 cases indicating a peak in biting time to occur before 22:00h (evening biting) and 9 cases indicating a peak in biting after 05:00h (morning biting) (Supplementary Table 1 and Supplementary Table 2). In about one third of those papers reporting evening or morning biting time, there was a suggestion of behavioural change in response to the use of LLINs. Further, a number of these papers reported measures of prevailing environmental temperature. In the majority (N = 21), the mean temperatures were 25°C or above (overall mean = 26.9°C), while the remainder (N = 12) had a lower mean of 21.4°C (Supplementary Table 1). There were significantly more cases of evening biting than morning biting overall (Chi-square test, *LR*-*χ*^2^ = 46.68, *df* = 1, *P* < 0.0001) regardless of temperature group (Fisher’s exact test, *P* = 0.171) (Supplementary Table 2).

### Effects of biting time and diurnal fluctuating temperature on vector competence

Having confirmed the potential for both morning and evening biting (including in major malaria vectors), we conducted a series of experiments to investigate whether biting time affected the potential for malaria vectors to become infected with the human malaria parasite, *Plasmodium falciparum*. Specifically, we aimed to determine the influence of both intrinsic (circadian) and extrinsic (diurnal temperature fluctuation) effects on measures of infection prevalence and intensity. We focused on the warmer temperature conditions as these were the most common in our literature search, are representative of high transmission settings, and are typical of conditions used in the majority of lab-based studies exploring human malaria-mosquito interactions. Accordingly, experiments were run on a 12:12h light:day cycle at either constant 27°C, or a more realistic mean temperature of 27°C with a Diurnal Temperature Range (DTR) of 10°C. Diurnal temperature ranges of 5-20°C are common across many malaria transmission settings^25, 33, 34^ and so DTR of 10°C is a representative intermediate value. Adult female mosquitoes of the African malaria vector, *A. gambiae*, were given infected blood meals at one of three times of the day to capture the range of potential feeding times from the evening through to the morning: 18:00h, 00:00h, or 06:00h. These times of day equate to Zeitgeber Times of ZT12, ZT18 and ZT0, respectively, where ZT0 refers to the beginning of the daylight cycle. For these time-of-day experiments, mosquitoes were maintained in separate incubators in which the timers were offset so that the actual feeds took place simultaneously using the same parasite culture, but the mosquitoes were at different points in their diel cycle. Note also that for the temperature fluctuation treatments, the mosquitoes were fed at 27°C, and then returned to their individual incubators to follow their particular diurnal thermal trajectories (Supplementary Fig. 1; see later discussion).

We found significant interactions between temperature and time-of-day on different measures of infection (oocyst intensity: Generalized Linear Mixed effects Model [GLMM], *F* = 17.36, *df* = 2, *P* < 0.0001; oocyst prevalence: GLMM, *F* = 18.64, *df* = 2, *P* < 0.0001; sporozoite prevalence: GLMM, *F* = 16.19, *df* = 2, *P* < 0.0001; Supplementary Table 3). Under the constant temperature regime there was no effect of time-of-day on oocyst intensity (i.e. number of oocyst in the midgut of infected mosquitoes), oocyst prevalence (i.e. proportion of mosquitoes infected), or sporozoite prevalence (i.e. proportion of mosquitoes with sporozoites in their salivary glands and hence, potentially infectious) (post-hoc contrasts, *P* > 0.05; Fig. 1). In contrast, under more realistic fluctuating temperatures, there was a significant effect of time-of-day on oocyst intensity, oocyst prevalence, and most importantly, sporozoite prevalence (post-hoc contrasts, *P* < 0.05; Fig. 1). Each of these infection measures was highest in mosquitoes fed at 18:00h (ZT12) and lowest in those fed at 06:00h (ZT0) (Fig. 1). For the 06:00h treatment, there was an approximate 98% reduction in sporozoite prevalence relative to the 18:00h treatment, with <1% of mosquitoes potentially able to transmit parasites. In addition, oocyst intensity and sporozoite prevalence was also lower in the 00:00h (ZT18) treatment compared to both the 18:00h (ZT12) treatment in the fluctuating temperature regime, and 00:00h (ZT18) in the constant temperature regime (post-hoc contrasts, *P* < 0.05; Fig. 1).

**Figure 1.**
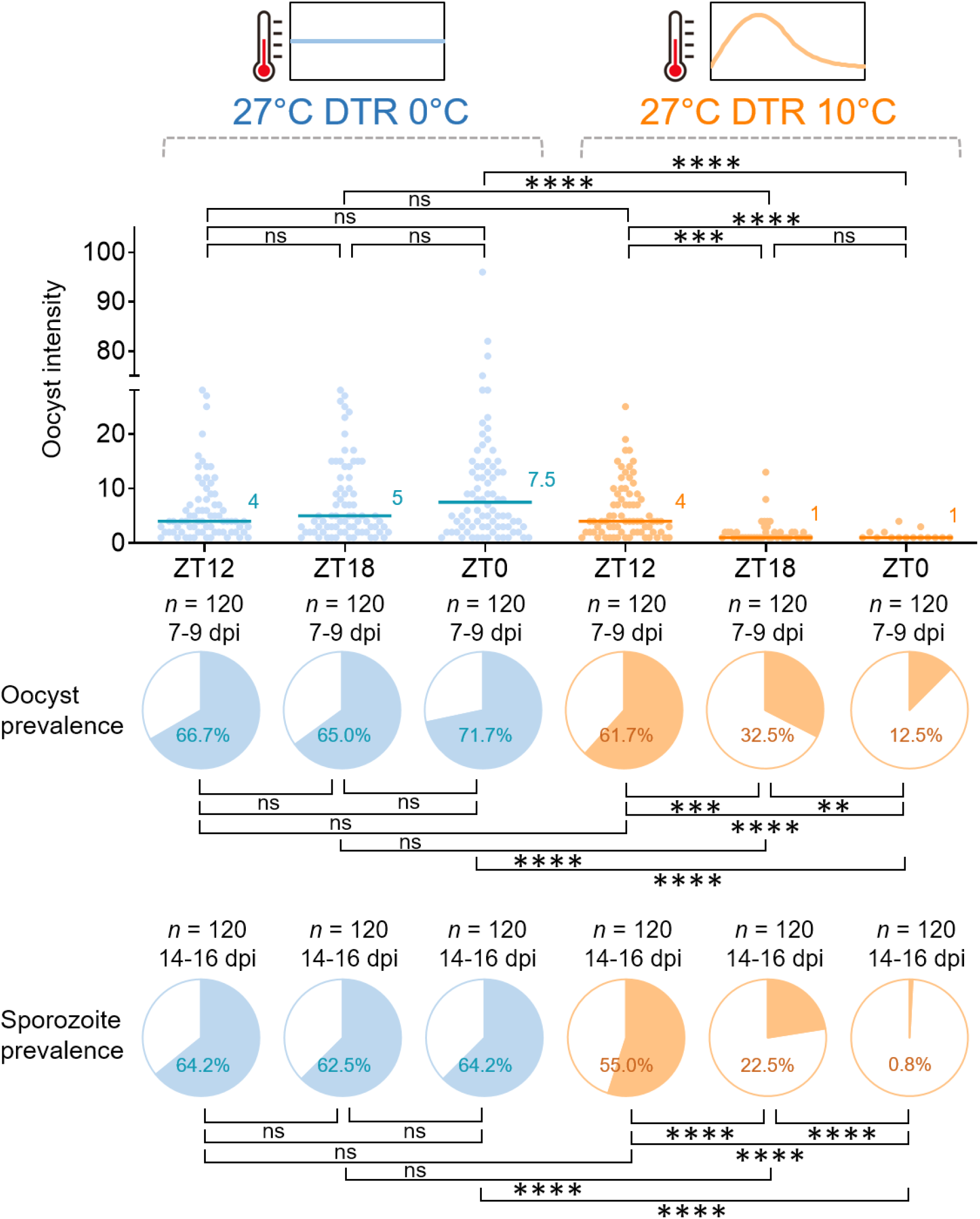
Effects of time-of-day of blood meal and diurnal temperature fluctuation on vector competence of *A. gambiae* mosquitoes infected with *P. falciparum* malaria. Mosquitoes were offered infected blood meals at a different time-of-day (18:00h [ZT12], 00:00h [ZT18], or 06:00h [ZT0]) and kept under either constant (i.e. 27°C with a Diurnal Temperature Range [DTR] of 0°C) or fluctuating (i.e. 27°C with a DTR of 10°C) temperature regimes. There is no effect of time-of-day of blood feeding on vector competence (oocyst or sporozoite prevalence) under constant temperature conditions but a significant increase in competence for mosquitoes feeding in the evening (18:00h; ZT12) and a significant reduction in competence for those feeding in the morning (06:00h;ZT0), relative to those feeding at midnight (00:00h; ZT18) under realistic fluctuating temperatures. The scatter plots show oocyst intensity, with the data points representing the number of oocysts found in infected individual mosquitoes, and the horizontal lines the median. The pie charts show oocyst or sporozoite prevalence calculated as the proportion of infected mosquitoes revealed by dissection of midguts and salivary glands, respectively. Asterisks indicate statistically significant differences between treatments (** *P* < 0.01, *** *P* < 0.001, **** *P* < 0.0001; ns, not significant at *P* = 0.05; *P*-values were Bonferroni corrected after pairwise comparisons). *n* indicates the number of mosquitoes sampled from four replicate containers of mosquitoes from two biologically replicated infection experiments. Forty mosquitoes were sampled daily from four replicate containers (10 per container) for dissecting midguts on 7-9 days post infection (dpi) or salivary glands on 14-16 dpi. Further details of the analysis are reported in Supplementary Table 3.

These results were corroborated for a second mosquito species, the Asian vector *A. stephensi*, in two separate infection experiments. In the first experiment, which followed the same experimental design described above, we found significant interaction between temperature and time-of-day on oocyst intensity (GLMM, *F* = 13.23, *df* = 2, *P* < 0.0001) and sporozoite prevalence (Generalized Linear Model [GLM], *LR*-*χ*^2^ = 14.08, *df* = 1, *P* < 0.001; Supplementary Table. 4). Consistent with the results for *A. gambiae*, under the constant temperature regime there was no effect of time-of-day on oocyst intensity, oocyst prevalence, or sporozoite prevalence (post-hoc contrasts, *P* > 0.05; Supplementary Fig. 2a). In contrast, under more realistic fluctuating temperatures, there was a significant effect of time-of-day on oocyst intensity, and more importantly, sporozoite prevalence (post-hoc contrasts, *P* < 0.05; Supplementary Fig. 2a). In the second experiment, we used a simplified design to provide a basic contrast between feeding in the evening (18:00h [ZT12]) vs morning (05:00h [ZT23]), under both constant and fluctuating temperatures. We found significant interactions between temperature and time-of-day on different measures of infection (oocyst intensity: GLM, *LR*-*χ*^2^ = 4.78, *df* = 1, *P* = 0.029; oocyst prevalence: GLM, *LR*-*χ*^2^ = 16.51, *df* = 1, *P* < 0.0001; sporozoite prevalence: GLM, *LR*-*χ*^2^ = 7.38, *df* = 1, *P* = 0.007; Supplementary Table 5). Again, oocyst intensity, and oocyst and sporozoite prevalence were not affected by time-of-day at constant 27°C (post-hoc contrasts, *P* > 0.05; Supplementary Fig. 2b), but all were significantly reduced when mosquitoes were fed in the morning under 27°C with a DTR of 10°C, compared with both the evening and morning feeds at constant temperatures (post-hoc contrasts, *P* < 0.05; Supplementary Fig. 2b).

### Effect of altered vector competence on malaria transmission potential

Our initial experiments suggest the potential for biting time to alter vector competence when daily temperatures fluctuate. To further explore the significance of these findings, we used a deterministic version of a transmission dynamics model of malaria^35–38^ to illustrate the potential public health implications of changes in vector competence in the context of LLIN use. First, we examined the effects of differences in vector competence alone on malaria prevalence, considering feeding distribution for an anthropophilic and anthropophagic vector where most bites happen at midnight and indoors (i.e. 70% at midnight and 30% in the evening and morning^39^), and illustrative scenarios where biting is skewed towards the evening (70% in the evening and 30% at midnight), or towards the morning (70% in the morning and 30% at midnight). Model predictions indicate no effect of biting time on malaria prevalence when all mosquitoes are equally competent (Fig. 2a and Supplementary Table 6). Similarly, when biting is centred around midnight there appears little effect of variation in vector competence (i.e. predicted malaria prevalence is almost identical whether competence differs between mosquitoes or not) (Fig. 2a and Supplementary Table 6). However, variation in competence leads to an increase in equilibrium infection prevalence if feeding is dominated by evening biting mosquitoes and a reduction in prevalence if feeding is dominated by morning biting (Fig. 2a and Supplementary Table 6). We next simulated the effects of LLINs assuming nets to be used by 50% of the population (approximating mean net use by children across sub-Saharan Africa^40^) and that contact rate with nets was the same for all mosquitoes regardless of biting time. LLINs reduced infection prevalence in all cases, but the relative efficacy is lower when biting is skewed towards the evening and greater when biting is skewed towards the morning, even when we assumed equivalent exposure to the LLINs for the different feeding behaviours (Fig. 2b and Supplementary Table 6). When we included the fact that evening and morning biters will likely experience reduced contact with LLINs (in the model, we halve the probability that biting takes place when people are in bed for the evening or morning biters), malaria prevalence increased overall, but the skew to evening biting resulted in the greatest prevalence and the lowest relative effectiveness of LLINs (Fig. 2c and Supplementary Table 6).

**Figure 2.**
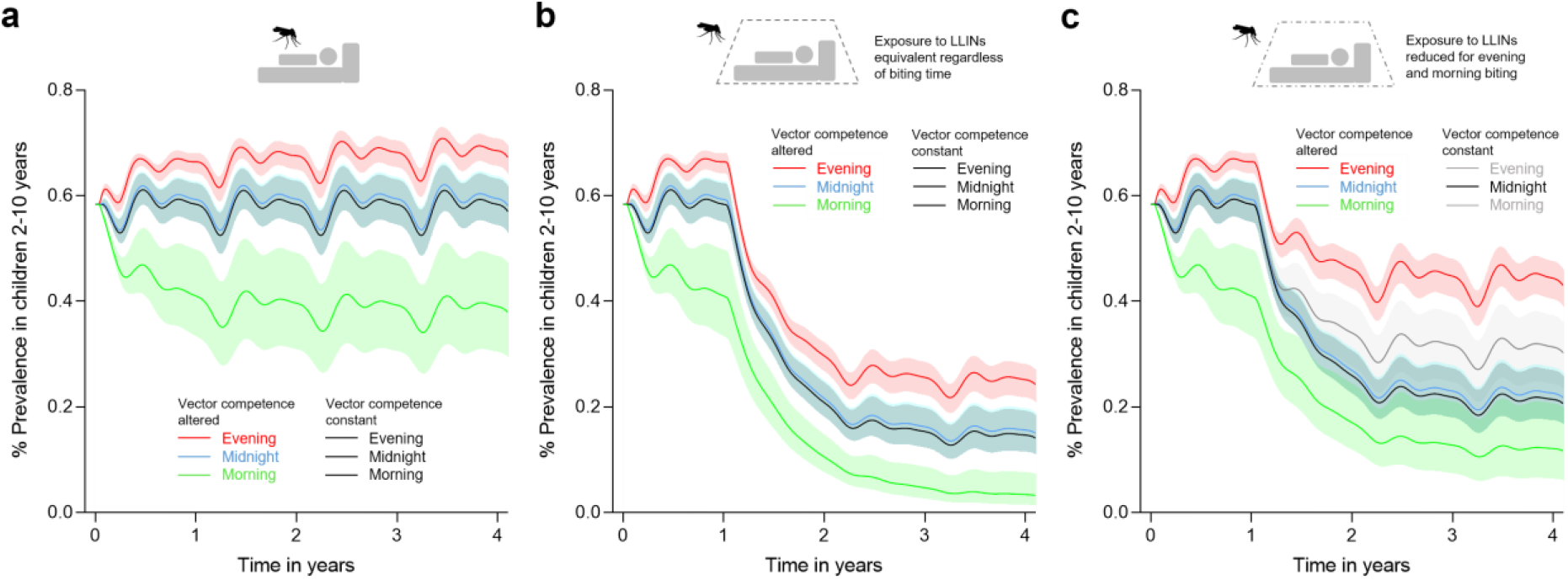
Model outputs illustrating potential epidemiological significance of altered vector competence. **a,** Effect of altered vector competence on malaria prevalence in children in a high transmission setting with mosquitoes biting predominantly in the evening (red line, run 2 and 5 in Supplementary Table 6), at midnight (blue line, run 1 and 4 in Supplementary Table 6) or in the morning (green line, run 3 and 6 in Supplementary Table 6) in the absence of bed nets (LLINs). In these and subsequent figures the solid lines represent the means and the matching coloured bands, the 95% confidence intervals. The black line shows the control scenarios where, in line with conventional assumptions, competence is the same for all mosquitoes (run 7-12 in Supplementary Table 6). In these cases of constant vector competence, prevalence is identical regardless of biting time. If we allow competence to vary in line with our empirical data (i.e. high for evening biters, intermediate for midnight biters and low for morning biters), there is little effect on prevalence if mosquitoes bite predominantly at midnight. However, variation in competence leads to increased infection prevalence when feeding patterns are skewed towards evening biting, and reduced prevalence when skewed towards morning biting. **b**, Impact of LLINs on malaria prevalence when mosquitoes bite predominantly in the evening, at midnight, or in the morning either with altered (evening = red line, midnight = blue line, morning = green line, run 1 – 3 in Supplementary Table 6) or constant (evening, midnight, and morning = black line, run 7 – 9 in Supplementary Table 6) vector competence, assuming all mosquitoes have equal probability of contacting an LLIN (i.e. the impact of LLINs on mosquito mortality and transmission potential does not vary with biting time). Under these assumptions, LLINs lead to reduced overall infection prevalence, but the efficacy of LLINs is less if biting is skewed towards the evening relative to midnight or morning biting, as evening biters have the greatest vector competence and hence, higher overall transmission potential. **c,** Impact of LLINs on malaria prevalence when mosquitoes bite predominantly in the evening, at midnight, or in the morning either with altered (evening = red line, midnight = blue line, morning = green line, run 4 – 6 in Supplementary Table 6) or constant (evening and morning = black line, midnight = grey line, run 10 – 12 in Supplementary Table 6) vector competence, but assuming that mosquitoes feeding in the evening or morning have reduced contact with LLINs (either because they feed outdoors or because people are less likely to be in bed and using nets at these times). Under these assumptions the relative efficacy of LLINs is reduced, but most markedly when feeding is dominated by evening biting mosquitoes with highest vector competence.

### Mechanistic effects of temperature fluctuation on vector competence

In order to better understand the influence of temperature fluctuation on vector competence, we conducted a series of experiments to determine the thermal sensitivity of malaria parasite establishment. The focus on initial parasite establishment is justified since it is only during the initial 24h following feeding that mosquitoes experience different conditions (i.e. they follow different short-term thermal trajectories as feeding occurs at different points on the fluctuating cycle) and conditions experienced in subsequent days are essentially identical. First, we examined the effects of absolute temperature by feeding *A. gambiae* and *A. stephensi* infected blood and maintaining them under constant temperatures of 27°C (control), 30°C, or 32°C, to test whether these higher temperatures were detrimental to parasite infections as temperature rise to >32°C during the day cycle of the 27°C DTR10°C regime. We observed a decline in overall oocyst intensity (GLM, *LR*-*χ*^2^ = 78.7, *df* = 1, *P* < 0.0001) and oocyst prevalence (GLM, *LR*-*χ*^2^ = 36.9, *df* = 1, *P* < 0.0001) at 30°C relative to 27°C for both mosquito species, while no oocyst infections were observed at 32°C (Fig. 3a and Supplementary Table 7). These data indicate that parasite establishment is constrained at temperatures that exceed 30°C. Next, we examined the importance of duration of exposure to high temperatures by varying the period of incubation at the permissive temperature of 27°C from 3 to 48h post blood meal, before moving mosquitoes to the more constraining temperature of 30°C, to test whether the earlier stage of parasite infection in particular is sensitive to high temperatures. In this case, overall infection levels were low because the parasite culture had unexpectedly low gametocytemia. Nonetheless, we found that incubating at 27°C for 12 to 24h led to a progressive recovery in oocyst intensity and oocyst prevalence rendering the infections statistically not different to those observed in a cohort maintained at 27°C (post-hoc contrasts, *P* > 0.05), while those mosquitoes transferred to 30°C before 12h showed no infections (Fig. 3b), indicating higher thermal sensitivity of early infection (i.e. < 12h post infection).

**Figure 3.**
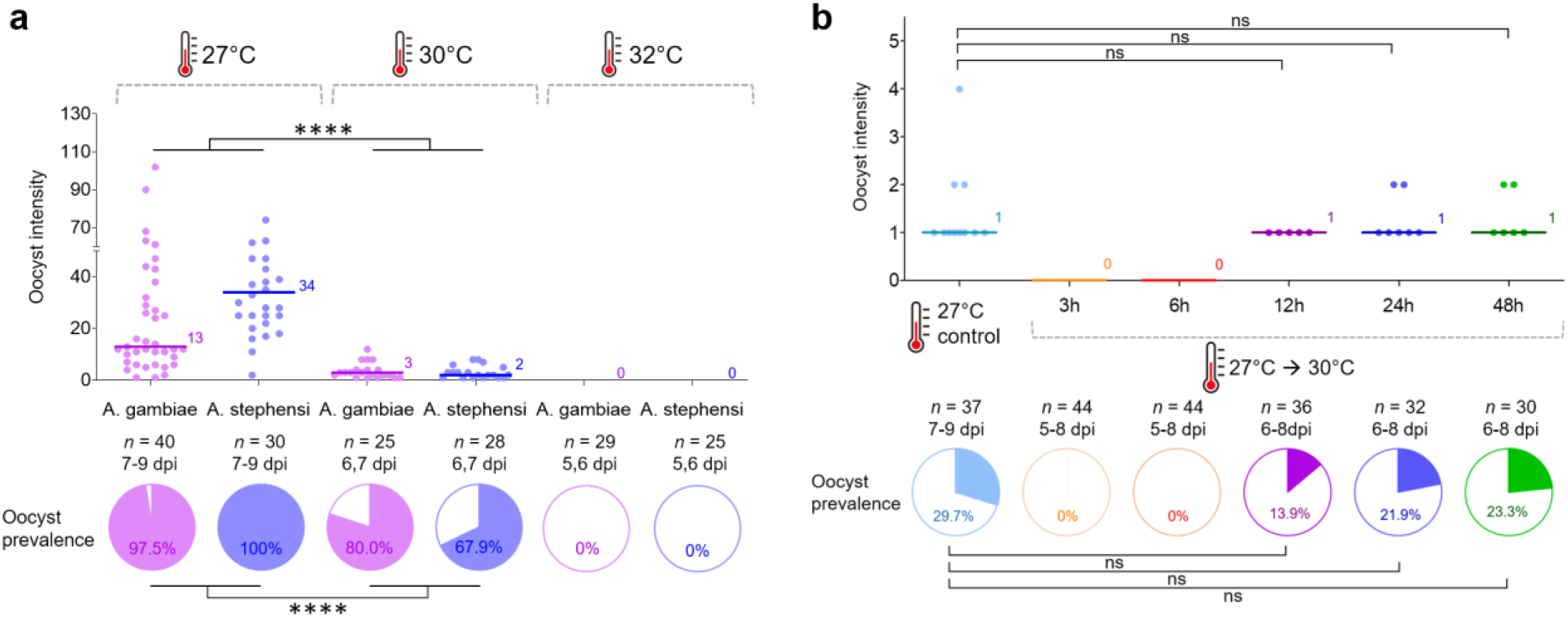
Effect of exposure to high temperatures on vector competence of *Anopheles* mosquitoes infected with *P. falciparum* malaria. **a**, *A. gambiae* and *A. stephensi* mosquitoes were kept at 27°C, 30°C, or 32°C following an infectious blood meal. The data indicate that exposure to constant 30°C is detrimental to parasite establishment for both *A. gambiae* and *A. stephensi,* while the infection is eliminated at 32°C. Results of analyses to examine the effects of temperature treatment and mosquito species on oocyst intensity or prevalence are reported in Supplementary Table 7. Asterisks indicate statistically significant differences between treatment groups (**** *P* < 0.0001). **b**, *A. stephensi* mosquitoes were incubated at 27°C for various periods of time ranging from 3 to 48h following an infectious blood meal, before being transferred to 30°C. Control mosquitoes were kept at 27°C throughout. These data indicate that the probability of parasite establishment in the mosquito increases as the time spent at a permissive temperature (27°C) increases, and that parasites are most sensitive to high temperatures during the first 12-24h following blood feeding. The control group was compared with each treatment group with > 0 infection using GLM with pairwise post-hoc contrasts followed Bonferroni corrections for *P*-values (ns, not significant at *P* = 0.05). For (**a)** and (**b)**, the scatter plots show oocyst intensity, with the data points representing the number of oocysts found in individual mosquitoes, and the horizontal lines the median. The pie charts show oocyst or sporozoite prevalence calculated as the proportion of infected mosquitoes revealed by dissection of midguts and salivary glands, respectively. *n* indicates the number of mosquitoes sampled per treatment group (dpi = days post infection).

An additional observation is that the effects of high temperature appear to vary to some extent with oocyst intensity (and so depend on the level of gametocytemia in the blood meal). For example, the data presented in Fig. 3a had the highest baseline intensities amongst our various experiments and in this case, reduction in oocyst prevalence at 30°C was not as high as when the baseline intensity was lower. To test the hypothesis that the negative effects of exposure to high temperature on parasite establishment depend on infection intensity, we fed *A. gambiae* blood meals containing four different dilutions (1, 1/2, 1/4, or 1/10) of gametocytes to generate a range of infection loads, and then kept them at 27°C or 30°C. At 27°C, the oocyst prevalence varied from 84 to 52% across the dilution treatments, with median oocyst intensities ranging from nine down to one per mosquito (Supplementary Fig. 3a). Incubation at 30°C reduced oocyst intensity and prevalence across the board (oocyst intensity: GLM, *LR*-*χ*^2^ = 5.96, *df* = 1, *P* = 0.015; oocyst prevalence: GLM, *LR*-*χ*^2^ = 138, *df* = 1, *P* < 0.0001; Supplementary Table 8). However, the per cent reduction in oocyst prevalence was 73% in the highest oocyst intensity treatment and increased up to 96% in the lowest intensity treatment (Supplementary Fig. 3a). Furthermore, when we plot per cent reduction in oocyst prevalence due to high temperature against mean number of oocysts per mosquito for each of our experiments, we find that the impact of temperature declines as intensity of infection increases (Supplementary Fig. 3b).

### Potential confounders

There are a number of potential confounders that could impact the robustness of our results. For example, we assume that in a fluctuating temperature environment, mosquitoes will generally track ambient temperature and not exhibit strong thermoregulatory behaviours that might limit exposure to the critical temperatures that impact parasite establishment. In order to investigate this, we adapted methods from a previous study^41^ to examine the thermal avoidance behaviour of *A. gambiae* following a blood meal at 06:00h that is either infected or uninfected. The approach exposes mosquitoes to temperatures that ramp gradually from 28 to >35°C and monitors the time point at which mosquitoes escape the warmed microenvironment (Fig. 4a). We found no evidence that mosquitoes were sensitive to temperatures of 30-32°C and only observed a thermal escape response as temperatures approached 35°C (Fig. 4b). There were no differences between infected and uninfected mosquitoes in escape response (Log-rank test; *χ*^2^ = 1.25, *df* = 1, *P* = 0.264) (dissection of mosquitoes from this experiment revealed oocyst prevalence of 60-75% with 5.5-9 median oocyst intensity).

**Figure 4.**
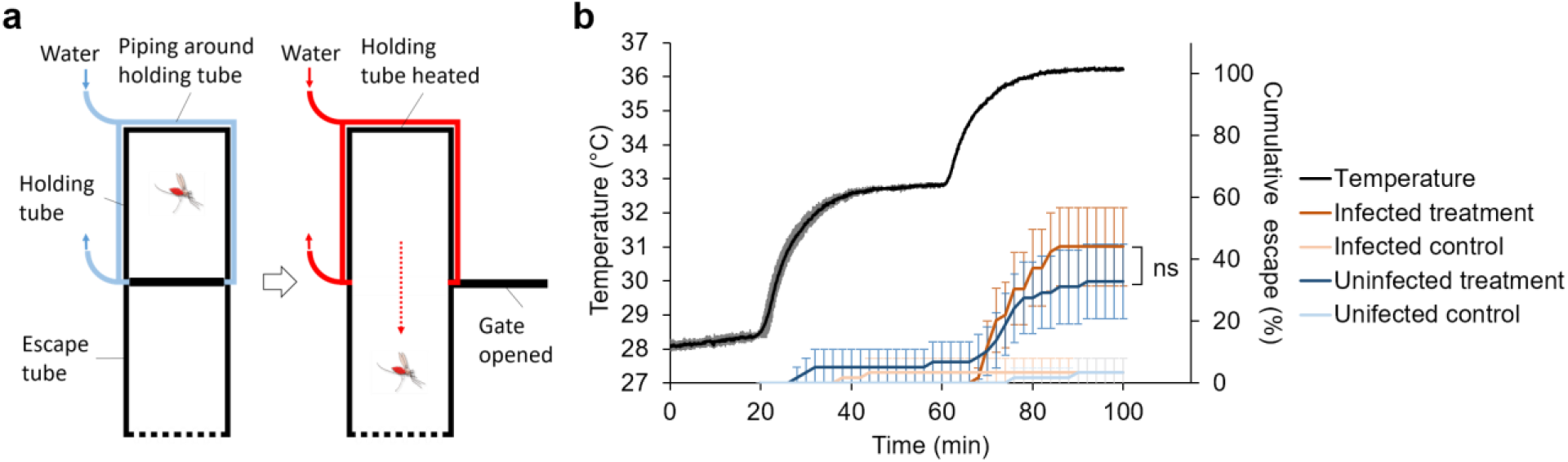
Behavioural assay to investigate thermal avoidance behaviour of *A. gambiae* mosquitoes following a blood meal. **a**, Diagram of the behavioural assay. The apparatus comprises two clear Perspex tubes joined by a sliding gate. One tube (the holding tube) is wrapped in plastic piping through which water is circulated. Infected or uninfected blood-fed mosquitoes (blood fed at 06:00h [ZT0]) are introduced into the holding tube and after a period of acclimation, the water is gradually heated from 28-36°C, and the sliding gate opened. The rate at which the mosquitoes leave the holding tube and enter the adjacent escape tube is recorded. For a control, the water is maintained at constant 28°C to measure baseline movement rates across the assay period for both infected and uninfected mosquitoes. **b**, Cumulative escape rate of infected and uninfected *A. gambiae* mosquitoes (error bars = 95% confidence intervals) in relation to temperature in the holding tube. The black line shows mean temperature with standard deviation (grey lines) in the holding tube in the ramping temperature treatment from three replicate runs. There were six replicates in total for each of the four mosquito groups (infected or uninfected, with either ramping temperature or constant temperature). The data reveal that mosquitoes were unresponsive to temperatures around 33°C, and only exhibited strong escape responses as temperatures was ramped up to 35°C and beyond. Control mosquitoes showed negligible movement across the assay period. These patterns were consistence whether mosquitoes had taken an infected or uninfected blood meal. Log-rank test was used to compare escape probability between the treatment groups (ns, not significant at *P* = 0.05).

Additionally, in our experiments the blood meal was administered at the mean temperature of 27°C before mosquitoes were returned to their respective temperature treatments. This was done to standardise blood feeding compliance and hence the proportion of mosquitoes acquiring parasites (note, blood feeding frequency exceeded 95% in all experiments). It is also technically challenging to blood-feed mosquitoes at different ambient temperatures for different temperature treatment groups using the same parasite culture at the same time. In reality, mosquitoes have to feed at the prevailing ambient temperatures. However, these prevailing temperatures for the different feeding times in the 27°C DTR 10°C regime vary from 22.6 to 28.5°C, so it is unlikely that these modest temperature differences would impact feeding compliance or efficiency, especially when the blood meal itself is at 37°C and this has a marked effect on mosquito body temperature during the feeding process^42, 43^. To provide some confirmation of this, we conducted a simple assay to compare the feeding efficiency of *A. gambiae* mosquitoes at 21 and 27°C. We found no effect of temperature or its interaction with time-of-day on feeding compliance (Temperature: GLMM, *F* = 3.05, *df* = 1, *P* = 0.131; Temperature × Time-of-day: GLMM, *F* = 3.98, *df* = 1, *P* = 0.080; Supplementary Table 9 and Supplementary Table 10). Furthermore, as part of a separate investigation, we have conducted an experiment in which *A. gambiae* adults were maintained at 21°C, fed at 27°C and then returned to 21°C to test whether transferring mosquitoes between different temperatures for blood feeding could affect vector competence. We found no difference in oocyst intensity, or oocyst or sporozoite prevalence between mosquitoes transferred between 21 and 27°C, and those maintained at 27°C throughout (post-hoc contrasts, *P* > 0.05; Supplementary Fig. 4).

We also examined whether transfer of mosquitoes at different times of day from their respective fluctuating temperatures affected subsequent blood meal size at the common feeding temperature of 27°C. Using fresh body weight of blood-fed mosquitoes as a proxy for blood meal size, we found no difference in body weight between temperature (time-of-day) groups for either *A. gambiae* or *A. stephensi* (GLMM, *F* = 0.46, *df* = 2, *P* = 0.635; Supplementary Fig. 5; Supplementary Table 11).

## Discussion

In the current study we used a combination of empirical and theoretical approaches to explore whether mosquitoes feeding at different times of day were equally likely to become infected with malaria parasites and hence contribute to transmission. The research was motivated by the fact that although most malaria mosquitoes tend to feed at night, the distribution in biting around the peak means that a proportion of bites also occur in the evening and the morning. Our analysis of the recent literature indicates that this crepuscular feeding is widespread (from our review around 50% of cases reporting hourly feeding behaviour indicated peak biting time either before 10pm or after 5am) and might possibly be increasing as a behavioural avoidance response to the use of insecticide treated bed nets. This suggestion is consistent with another recent systematic review, which indicated that on average only 79% of bites by the major malaria vectors in Africa occur during the time when people are in bed, an estimate substantially lower than previous predictions^16^. Note also that there are very broad confidence intervals around this estimate, with 95 percentiles ranging from 33.9 to 97.2% for bites received when people are in bed, depending on vector species and location^16^.

How such feeding behavior influences transmission depends, in part, on whether biting time affects the capacity of mosquitoes to acquire and successfully incubate the malaria parasite. From a range of laboratory infection studies, we show that vector competence varies substantially depending on whether mosquitoes feed in the evening, at midnight, or in the morning. This variation does not appear to be driven by circadian rhythm of the mosquitoes but rather, an interaction with daily temperature variation. More specifically, time-of-day of feeding had no significant effect on the proportion of mosquitoes that successfully developed parasites through to sporozoite stage when mosquitoes were maintained at constant 27°C. However, when mosquitoes were maintained under conditions representing more realistic diurnal temperature variation (i.e. 27°C±5°C) there was significant variation in vector competence, with approximately 55 and 88% of evening biters, 26 and 65% of midnight biters, and 0.8 and 13% of morning biters positive for sporozoites for *A. gambiae* and *A. stephensi*, respectively (Fig. 1 and Supplementary Fig. 2a). Consistent with some earlier work^29, 30^, our additional experiments suggest that this pattern results from transient exposure to temperatures >30°C reducing vector competence via a negative effect on the initial stages of parasite development. Importantly, mosquitoes feeding in the morning (i.e. 06:00h [ZT0]) have only 4h before temperatures exceed 30°C under a fluctuating temperature regime, while those that feed at midnight or in the evening (i.e. 00:00h [ZT18] or 18:00h [ZT12]) have 10h and 16h at permissive temperatures, respectively (see Supplementary Fig. 6). As the duration of permissive temperatures increases, so does the probability of parasite establishment.

Our illustrative modelling analysis indicates that the differences in vector competence associated with biting time could have important implications for malaria burden (Fig. 2). In the absence of LLINs, the variation in vector competence we observe in our empirical studies leads to increased infection prevalence in the human population when feeding patterns are skewed towards evening biting, and reduced prevalence when skewed towards morning biting. When biting is distributed symmetrically around midnight the model suggests negligible effect of variation in competence on prevalence, relative to predictions based on the standard assumption that all mosquitoes have equal competence. However, this does not mean that variation in competence is unimportant but rather, that the increased transmission potential of mosquitoes biting in the evening is more or less counterbalanced by reduced transmission potential of mosquitoes biting in the morning. LLINs reduce overall infection prevalence, but the impact of LLINs is less if biting is skewed towards the evening relative to midnight or morning biting, as evening biters have the greatest vector competence and hence, higher overall transmission. If we further assume that evening or morning biting mosquitoes escape contact with bednets because people are unlikely to be in bed and protected by LLINs at these times, the relative efficacy of LLINs is reduced, even if mosquitoes have equivalent competence (comparing the grey lines with the black lines in Fig 2c provides an illustration of the impact of behavioural resistance with constant competence). If we include the additional effect of variable vector competence, the decline in relative efficacy of LLINs is more modest for morning biters but greater for evening biters. The reason is that if mosquitoes feed in the morning, the reduced competence of the mosquitoes could compensate for the lower contact rate with LLINs. In the case of morning feeding being a consequence of behavioural resistance, such an effect would represent an unexpected positive side effect of selection on mosquito life history^44^. On the other hand, if mosquitoes feed in the early evening, then not only will LLIN contact rate tend to be reduced, but the mosquitoes could be even more efficient vectors, exacerbating the epidemiological consequences of residual transmission and/or behavioural resistance.

The exact mechanisms underlying the transient thermal sensitivity of parasite establishment remain unclear. There could be direct negative effects of temperature on parasite biology and/or indirect effects mediated via the mosquito. Previous research has suggested that an increased blood digestion rate at higher temperatures could increase the quantity of midgut proteases, potentially reducing ookinete density in the mosquito midgut^30^. Given the importance of elements of the innate immune response and certain components of the midgut microbiome in determining susceptibility to infection, it is possible that these factors could also interact with temperature^45, 46^.

Our results appear robust to mosquito behaviour as our thermal escape response assay indicated that mosquitoes are behaviourally unresponsive to temperatures that are critically damaging to malaria parasite establishment. The limited behavioural response of adults to temperatures of around 32°C is similar to that reported previously^41^. Moreover, studies comparing the effects of temperature extremes on *Anopheles* mosquitoes indicate that long-standing laboratory colonies are sufficiently similar in thermal tolerance to field-collected mosquitoes to provide reasonable surrogates of wild populations^47^. Further, our feeding compliance and blood meal volume assays suggest that transferring mosquitoes between temperatures for feeding in our main experiments, likely had little confounding effect.

We acknowledge that our study used standard laboratory mosquito and parasite strains, and it is possible that in field settings, local adaptation could yield different patterns of thermal sensitivity for parasites in wild type mosquitoes^48^. Previous studies do indicate that infection with *P. falciparum* is possible above 32°C^49, 50^, and there is a suggestion that naturally circulating parasites might exhibit higher thermal tolerance than standard lab strains^51^. For example, one study using parasite populations from 30 naturally infected children in Kenya found that parasites established in mosquitoes following blood feeds from 50% of the carriers (i.e. blood from 15 of the 30 gametocyte positive children yielded mosquito infections) when mosquitoes were maintained at 27°C, but this fell to 30% (i.e. mosquito infections from blood of 8/27 of the children) when mosquitoes were maintained at 32°C ^51^. For those feeds that yielded infections at both temperatures, the mean percentage of mosquitoes infected at oocyst stage was 31% at 27°C and 17% at 32°C. These reductions in frequency of infection and infection prevalence are less extreme than our data might predict but still indicate a marked impact of temperature. Whether the differences between studies result from variation in parasite thermal sensitivity between strains, or other factors, is not known. Our data, together with those of Bradley *et al.*^52^ and Pathak *et al*. ^53^, suggest variation in gametocyte densities between feeds/hosts could mediate the effects of temperature on parasite establishment (i.e. if infection is partly a numbers game, then low gametocyte densities might result in even lower probability of a successful infection under thermally constraining conditions). There might also be circadian patterns in the developmental rhythm of parasites^54^ and gametocyte infectiousness^23^. Our experiments used cultured parasites and we found little evidence for circadian effects in the mosquito in the absence of temperature fluctuation. Recent work on rodent malaria, however, indicates that gametocytes are less infective in the day than at night, but this reduced infectivity is partly offset by mosquitoes being more susceptible to infection when they feed during the day^55^ (though it should be noted that neither the mosquito or the rodent species used in these latter studies is the natural host, and the infection experiments were conducted under constant temperatures). Studies on *P. falciparum* in the field provide mixed results; some research indicates no difference in infectiousness or density of gametocytes between day (16:00h) and night (23:00h)^56^, while other research suggests a diel cycle in gametocyte density with the highest density in the early evening (17:30h) and the lowest in the morning (05:30h) ^57^, which would likely exacerbate the effects we report.

In addition to potential biological differences between systems (both lab vs. field, and field vs. field), how the time-of-day effects impact malaria transmission intensity in the field will likely vary with prevailing temperatures. If either the mean temperatures or the extent of daily temperature variation limit exposure to temperatures above 30°C, there might be little impact of biting time. Whether biting time affects competence in conditions representative of the lower temperature environments we identified in the literature review is the subject of ongoing research. However, there are extensive areas of malaria transmission in Africa where peak daily temperatures exceed 30°C^25–28^. Furthermore, interactions with other traits could influence the net impact on transmission. For example, it is generally assumed that malaria vectors feed at night to exploit sleeping hosts and reduce biting-related mortality^15^. The extent to which feeding earlier in the evening increases mortality rate or otherwise influences mosquito-to-human transmission and thus vectorial capacity overall, is unknown. Further mathematical modelling work is needed to better understand the full implications of the difference in human-to-mosquito transmission, though it will be impeded by a general lack of knowledge of mosquito behaviour and transmission ecology^58–60^.

All these factors caution against over-extrapolation of our results and point to the need to extend research to field settings to validate our findings using natural mosquito-parasite pairings. Nonetheless, the high thermal sensitivity of the early stages of malaria parasite infection is widely observed in diverse systems, including human (*P. falciparum* and *P. vivax*), rodent (*P. chabaudi* and *P. berghei*), and avian malaria (*P. relictum*)^29, 30, 33, 61–63^, so there is little reason to think the qualitative effects we report are unique to our experimental system. As such, we believe our empirical and theoretical findings could have significant implications for basic understanding of malaria transmission ecology since they suggest that not all mosquito bites are equivalent and that evening feeding might contribute disproportionately to vectorial capacity. There is significant interest in how aspects of the innate immune system^64–66^, or factors such as the midgut microbiome^67–69^, can impact the capacity of mosquitoes to transmit malaria parasites. In the context of this research, it is noteworthy that ecological factors like daily variation in temperature and biting time can interact to render the same mosquitoes either highly susceptible, or essentially refractory. These results are not simply of academic interest as they add important ecological complexity to understanding the potential significance of residual transmission and behavioural resistance.

## Materials and Methods

### Characterization of biting behaviour in *Anopheles* mosquitoes in the literature

We used eight combinatorial search terms composed of ‘biting’, either of ‘malaria’ or ‘Anopheles’, and one of ‘nets’, ‘bednets’, ‘ITNs’, or ‘LLINs’ in PubMed for identifying literature that provided hourly biting time data (18:00-06:00h) generated by human landing catch methods or human baited bed net traps. Publication year was limited to 2000-2017 considering a marked increase for the malaria control efforts in sub-Saharan Africa since 2000^1^. Conventional peak biting time of *Anopheles* mosquitoes is generally known to occur between 00:00-04:00h^15, 32^, and studies have shown majority of people go to bed at 21:00-22:00h and get out of bed at 05:00-06:00h^7, 70–75^. Accordingly, we considered cases of peak biting time before 22:00h (i.e. evening biting) or after 05:00h (i.e. morning biting) to be consistent with behavioural change. A “case” was determined as a mosquito species or species complex for which biting activity had been determined in a given paper.

Temperature data for the studies were either provided directly in the source literature or, if not presented, monthly mean temperature was estimated for the time (study periods) and location (regional estimates of study sites) of the study using Global Surface Summary of the Day (GSOD) provided by National Oceanic and Atmospheric Administration, Department of Commerce (https://data.noaa.gov/dataset/dataset/global-surface-summary-of-the-day-gsod), or Climte-Data.org (https://en.climate-data.org). Temperature measures were categorized into high (25°C or above) and low (< 25°C) based on a recent study determining the optimal temperature for malaria transmission as 25°C^76^, and a mean temperature was determined for each group.

### Mosquitoes

*Anopheles gambiae* (G3, NIH) and *A. stephensi* (Liston, Walter Reed Army Institute of Research) mosquitoes were used throughout the experiments. Mosquitoes were reared under standard insectary conditions at 27°C±0.5°C, 80%±5% relative humidity, and a 12h:12h light-dark photo-period. Larval density and amount of larval food (ground TetraFin^TM^; Tetra, Blacksburg, VA) were standardised to ensure uniform adult size. Adults were maintained on 10% glucose solution supplemented with 0.05% para-aminobenzoic acid (PABA). For the infectious feeds, 5-6-day-old female mosquitoes were randomly aspirated into cardboard cup containers that are covered with netting, and starved for approximately 6 hours before infectious feed. Individual containers contained 120-150 mosquitoes.

### General procedures for mosquito transmission studies

*In vitro* cultured *Plasmodium falciparum* (NF54 isolate, MR4) was provided by the Parasitology Core Lab (http://www.parasitecore.org/) at John’s Hopkins University. Gametocyte culture in stage four to five (day 14 after gametocyte initiation) was transported overnight to Penn State in a sterile 50ml falcon tube filled with fresh culture media. The culture tube was packaged in a Styrofoam box with heating pads to keep the temperature at approximately 37°C during transport. Gametocyte-infected erythrocytes were provided with fresh culture media on the day of arrival, and were maintained > 24 hours before the infectious feed to allow additional maturation of gametocytes.

Mosquitoes were fed on day 16 post gametocyte initiation. The proportion of erythrocytes infected with mature gametocytes (i.e. gametocytemia) generally ranged between 1-3% in the culture. An infectious blood meal was prepared by mixing gametocyte infected erythrocytes with fresh human serum and erythrocytes at 40% haematocrit on the day of blood feeding as previously described^77^. Gametocytemia in the blood meal was adjusted so that mosquitoes were infected at realistic infection intensities (e.g., see Supplementary Fig. 3, and Bradly *et al*.^52^).

All infectious feeds were conducted in a walk-in environment controlled chamber. Glass bell jars were uniformly covered with Parafilm to serve as membrane feeders and heated to 37°C with continuously circulating water as previously described^77^. An appropriate amount of infectious blood (1-2 ml depending on the size of experiment but consistently the same amount within an experiment) was pipetted into each bell jar. Containers of mosquitoes were randomly allocated to bell jars to minimize any effect of position or feeder. Mosquitoes were fed for 20 min at 27°C after acclimating at 27°C for an hour, and > 95% mosquitoes were fully engorged in all infectious feeds. Immediately after blood feeding, mosquitoes were placed into incubators (Percival Scientific Inc., Perry, Iowa) with appropriate temperature treatment conditions (90%±5% relative humidity, and 12h:12h light-dark photo-period) and provided daily with fresh 10% glucose solution supplemented with 0.05% PABA. Mosquitoes were transferred and fed under red light as appropriate to maintain light:dark cycles.

To determine vector competence, mosquitoes were randomly collected by aspirating into 95% ethanol, and midguts and salivary glands were dissected in 1× phosphate-buffered saline solution under a standard dissecting scope. Presence or absence of parasite infection was determined by examining midguts and salivary glands, and oocysts in midguts were counted, using a compound microscope. To ensure correct scoring, oocysts and sporozoites were inspected under 40× magnification and cross-checked by a second person. Oocyst or sporozoite prevalence was calculated as the total number of infected mosquitoes divided by the total number of dissected mosquitoes by combining dissection data from given dissection days and replicated containers of mosquitoes for each treatment.

### Experimental design for mosquito transmission studies

#### Effects of biting time and diurnal fluctuating temperature on vector competence

For experiments examining the effect of time-of-day of blood meal and diurnal temperature fluctuation on vector competence, *Anopheles* mosquitoes were infected at different times of day and maintained at 27°C with a Diurnal Temperature Range of zero (i.e. DTR 0°C) or with a DTR of 10°C (i.e. 27°C±5°C; Supplementary Fig. 6). The Parton-Logan model was used for the fluctuating temperature regime that follows a sinusoidal progression and an exponential decay for the day and night cycle, respectively^33, 78^. The air temperature of incubator (Percival Scientific Inc., Perry, Iowa) was monitored closely using HOBO data loggers (Onset Computer Corporation, Bourne, MA) at 5 min intervals, and the accuracy of temperature was maintained with the error range of ± 0.5°C. Prior to infections, pupae were collected and placed into separate incubators in which the clocks were offset so that adult mosquitoes emerged into environments that were staggered in terms of time-of-day. This enabled us to do the infectious feeds simultaneously using the same parasite culture, but with the mosquitoes at different points in their diel cycle (see Supplementary Fig. 1). *Anopheles* mosquitoes were provided with infectious blood meals in two containers of mosquitoes (150 each unless otherwise specified) at 18:00h (ZT12), 00:00h (ZT18), or 06:00h (ZT0) and maintained at either 27°C with DTR 0°C or DTR 10°C (i.e. two replicates per treatment group). For dissections, twenty mosquitoes were sampled daily (10 per container) on 7, 8, and 9 days post infection (dpi) for oocysts and 14, 15, and 16 dpi for sporozoites. Oocyst intensity, or oocyst or sporozoite prevalence were determined using dissection data from the three days (sample size of 60 per treatment). We repeated the experiment two times for *A. gambiae* and one time with *A. stephensi*, each with different batches of parasite culture and mosquitoes. A further independent experiment was conducted with *A. stephensi* in which approximately 150 *A. stephensi* mosquitoes were fed in a container at 18:00h (ZT12) or 23:00h (ZT23) and maintained at either 27°C with DTR 0°C or DTR 10°C. Approximately 10 mosquitoes were sampled daily on 8-10 dpi for dissecting midguts to determine oocyst intensity or prevalence, and on 13, 14, and 16 dpi for dissecting salivary glands to determine sporozoite prevalence.

#### Effect of temperature variation on vector competence

Effects of temperature treatment, mosquito species and/or gametocytemia on vector competence were examined in a series of infection experiments. For general procedures, approximately 120 mosquitoes were fed in a container (unless otherwise specified) with *P. falciparum* infected blood meals, and maintained at appropriate temperature conditions for each experiment. Approximately 10-15 mosquitoes were collected daily for generally 2-3 days to dissect midguts or salivary glands, unless otherwise specified. Dissection days were determined by Detinova’s parasite growth model^79^ and data from pilot tests (data not shown) to ensure we sampled when infection prevalence was at a maximum depending on temperature treatments. For measures of vector competence, oocyst intensity, or oocyst or sporozoite prevalence were determined by combining data among dissection days. A separate batch of parasite culture was used for each experiment, and mosquitoes were fed around 18:00h (ZT12) to standardise time-of-day of blood feeding, unless otherwise specified.

In the first experiment, infected *A. gambiae* and *A. stephensi* mosquitoes were maintained at 27°C, 30°C, or 32°C to examine the effect of high temperature on vector competence. In the second experiment, to examine the effect of high temperature on early parasite infection, *A. stephensi* mosquitoes were incubated at 27°C for 3h, 6h, 12h, 24h, or 48h before moving them to 30°C. As a control group, infected mosquitoes were maintained at 27°C. In the third experiment, to examine the effects of gametocytemia and temperature interaction on vector competence, *A. stephensi* mosquitoes were fed blood meals with varying gametocytemia dilutions (1, 1/2, 1/4, or 1/10) and maintained at 27°C or 30°C. An infectious blood meal was prepared as described above, and serially diluted to generate blood meals with lower gametocytemia while maintaining 40% haematocrit. In the fourth experiment, 240 *A. gambiae* mosquitoes were fed in two containers (120 each) and kept at 21°C to examine the effect of transferring mosquitoes between different temperatures. Prior to the infection, pupae were collected and placed into the incubator at 21°C. As a control, mosquitoes were kept at 27°C throughout. Control and treatment mosquitoes were fed at 27°C (at 00:00h [ZT18]).

#### Feeding compliance and blood meal size

To determine the effect of different temperatures on blood feeding compliance, we compared feeding rates of *A. gambiae* maintained at 21°C DTR 0°C with the 27°C DTR 0°C control (data from Fig. 1, 2^nd^ feed). Mosquitoes were reared as described above. Mosquitoes were provided with infectious blood meals in two containers (120 each) at 18:00h (ZT12), 00:00h (ZT18), or 06:00h (ZT0). Blood feeding compliance was measured by scoring the proportion of unfed mosquitoes.

To explore whether transfer of mosquitoes from different points on the fluctuating cycle (i.e. 18:00h [ZT12], 00:00h [ZT18], 06:00h [ZT0] in the 27°C DTR 10°C temperature regime) affected subsequent blood meal size of mosquitoes feeding at 27°C, we compared the body weight of blood-fed mosquitoes as a proxy for blood meal size. Mosquitoes were reared following the same protocol for the time-of-day and fluctuating temperature experiment described above. The blood meal was prepared using the same method used for the infectious feeds, except we used uninfected blood on this occasion. After starving for 6h prior to blood feeding by removing the sugar source, 5∼6 day old *A. gambiae* and *A. stephensi* female mosquitoes were blood fed for 20 min at 27°C in two containers (30 each) per each time-of-day treatment with 1h acclimation at 27°C. One hour post blood feeding, blood-fed mosquitoes were killed by freezing at −20°C for 30 min, and unfed mosquitoes were discarded. Twenty mosquito samples were randomly selected from each container to measure the whole body weight of individual mosquitoes (i.e. 40 sample size per treatment group per species), using an analytical balance with the accuracy of ±0.1mg (MS104S; Mettler Toledo, Columbus, OH).

### Thermal avoidance assay

*A. gambiae* mosquitoes were collected at pupal stage and adapted for > 5 days at 27°C DTR 10°C until blood feeding. Mosquitoes were fed with either *P. falciparum* infected or uninfected blood meals at 06:00h (ZT0) as described above, and maintained at 27°C until used for the behavioural assay. Three containers of mosquitoes were fed (100 mosquitoes per container) for the infected or uninfected groups, and mosquitoes from a container from each group were used for each round of assay. Infected and uninfected blood meals were prepared as described above, but gametocyte infected-erythrocytes were replaced with uninfected erythrocytes in the uninfected blood meal. The behavioural assay was conducted in an environmental chamber at 27°C±0.5°C with 80%±5% relative humidity using WHO insecticide bioassay tubes as described previously^41^ (Supplementary Fig. 7). One side of the tube (the holding tube) was wrapped with plastic tubing with continuously circulating water heated by a water bath (WB05; PolyScience Inc., Niles, IL) to control the inner surface temperature of holding tube, while the temperature of escape tube was maintained at 28°C±0.5°C. Ten mosquitoes fully engorged with either infected or uninfected blood meals were introduced into a holding tube and acclimated at 28°C±0.5°C. The assay tubes were used in rotation by mosquito groups fed with either infected or uninfected blood meals within and between the assays. The gate between the holding and escape tubes was opened after 20 min of acclimation, and mosquitoes could then choose to move to the escape tube. The number of mosquitoes in the escape tube was recorded every 2 min. No mosquitoes in the escape tube returned ever to the holding tube during the entire assay period. The temperature of water bath was set to 32.6°C at the time of gate was opened, which was equivalent to the maximum temperature in 27°C DTR 10°C treatment. The surface temperature of holding tube increased at the rate of approximately 0.23°C/min over 20 min and was maintained at 32.6°C±0.5°C for an additional 20 min. The temperature of the water bath was then set to 36°C to further examine the thermal behaviour. The surface temperature of holding tube increased at the rate of approximately 0.17°C/min over 20 min, and was maintained at 36°C±0.5°C for additional 20 min. The rates of temperature increase were comparable to that of Kirby and Lindsay^41^. The temperature of the two holding tubes in the treatment group was recorded at 5 sec intervals using thermocouple data loggers (SL500; MicroDAQ.com, Ltd., Contoocook, NH) throughout the experiments. Baseline activity of mosquitoes were monitored as a control by keeping the temperature of the holding and escape tubes at 28°C±5°C throughout the experiment and otherwise following the same methods as for the treatment group. A total of eight assay tubes were used for running the control and treatment groups (four each) at the same time, with two replicates for the mosquito group fed with infected or uninfected blood meals. Three rounds of assay were conducted between 4-10 hours post infection totalling six replicates for each mosquito group (see Supplementary Fig. 7 for experimental setup). Oocyst prevalence and intensity were determined on 8 dpi in total cohort of 60 mosquitoes (20 per container) fed with the same infectious blood meal and kept at 27°C.

### Transmission dynamics model

A deterministic version of a transmission dynamics model of malaria^35–38^ was adjusted and used to explore the potential public health implications of a theoretical change in mosquito infectivity driven by the timing of mosquito bites. The transmission model mechanistically tracks *P. falciparum* infection in people and mosquitoes. Susceptible people are exposed to infectious mosquito bites at a rate dependent on local mosquito density and infectivity. Mosquito dynamics describe the effects of mosquito control and the resulting decline in egg laying^36^. Adult mosquitoes can be either susceptible to malaria, infected, or infectious to people.

#### Model adjustment

Susceptible (S_V_), exposed (E_V_) and infective (I_V_) mosquitoes are present, and the dynamics of these are expressed as:

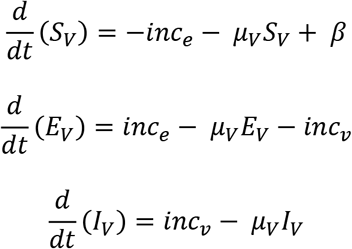

Where mosquito pupae emerge as adults at a rate *β* that is density dependent and die at a constant rate *μ_V_* which is assumed to be independent of infection status. The rate of exposure to infection with *Plasmodium falciparum* (*inc_e_*) is dependent on the force of infections (*FOI_V_*) such that:

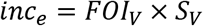

The Force of Infection is the product of: i) the sum of infectivity from all infective people in the population, ii) the biting rate, which is allowed to vary such that some people could be bitten more than others; iii) the biting rates of mosquitoes, which is dependent on vector intervention categories (bed nets and/or indoor residual spraying may be used), and; iv) the age-specific force of infection, which is normalized so that people of different ages could contribute differently to transmission.

The rate that mosquitoes then became infectious (*inc_v_*) is dependent on the proportion of mosquitoes that survived long enough to transmit infection onward. The extrinsic incubation period (EIP) is the number of days needed for sporozoites to be found in the salivary glands. The current model has a constant EIP of 11.5 days (to match experimental data).

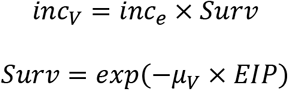

Based on the experimental observations, the transmission model was adjusted so that mosquitoes that exhibit either evening (*Mosq_EV_*), midnight (*Mosq_MD_*) or morning (*Mosq_MN_*) biting behaviour had different rates of becoming infective (*inc_V_*).

The differential equations describing the infection status of adult mosquitoes is adjusted such that:

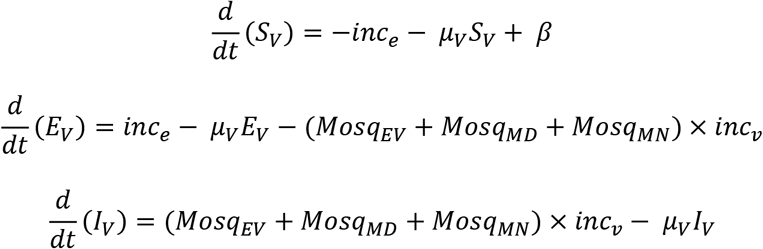

The sum proportions of mosquitoes in the different feeding classes equals 1 but the conversion to sporozoite positive status is different for each group. Our empirical study showed that about 55% of mosquitoes that bite in the evening became infectious (i.e. sporozoite positive), compared to 22.5% that bite at midnight and 0.83% of those that bite in the morning. The conversion term, *tob*:

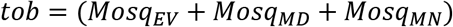

was parameterized to sum to 1 when all mosquitoes are feeding at midnight. The proportion of mosquitoes was adjusted given the ratio of 55.0%:22.5%:0.83% for vector competence in evening, midnight and morning biters respectively, such that:

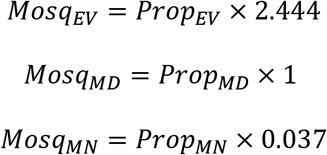

For baseline (control) scenarios representing constant vector competence with respect to biting time, the proportion of mosquitoes was adjusted given the ratio of 22.5%:22.5%:22.5% for vector competence of evening, midnight and morning biters, respectively (assuming all equals to midnight biting), such that:

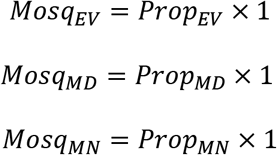

To estimate 95% confidence intervals (CIs) for prevalence data, exact Clopper-Pearson 95% CI was estimated from the empirical vector competence data of 55% (45.7-64.1%), 22.5% (15.4-31.0%), and 0.83% (0.02-4.7%) for evening, midnight and morning biting, respectively. These CI values were then divided by the standard prevalence of 22.5% to calculate equivalent parameter ratios of 2.444 (2.029-2.848), 1 (0.684-1.379) and 0.037 (0.001-0.203) for evening, midnight and morning biting, respectively.

#### Mosquito biting patterns

Evidence suggests that most mosquitoes actively search for blood-meals in the middle of the night and less so either in the evening or morning^71, 74, 80, 81^. To reflect this, we considered a ‘status quo’ scenario that examines the proportion of mosquitoes feeding during the evening and morning to be 0.15 each whilst the proportion feeding at midnight is 0.7 (as per Supplementary Table 6; runs 1, 4, 7, and 10). We then explored what would happen if mosquito feeding patterns shifted toward evening (Supplementary Table 6; runs 2, 5, 8, and 11) or morning (Supplementary Table 6, runs 3, 6, 9, and 12).

#### Contact with bed nets

The degree of protection that a bed net can elicit depends on the proportion of bites received while a person is protected. Therefore, in the transmission model, the impact of bed nets is determined by the proportion of bites that happen when a person is in bed (Φ_B_). Bed nets are modelled to impact the probability of mosquito from species *i* successfully biting (w_i_), and the probability of repellence (a mosquito is reflected away by the intervention before biting) (z_i_) following (1):

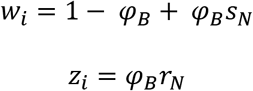

Mosquitoes that successfully feed (*s_N_*), die (*d_N_*) or repeat a feeding attempt (*r_N_*) in the presence of a bed net relative to the absence of a bed net were estimated using data from experimental hut trials that examined the entomological impact of LLINs^2, 35^. People are usually not in bed at 18:00h and start getting up before 06:00h^82, 83^, thus the probability of LLIN contact varied by mosquito biting time (in reality, these proportions may vary night to night or person to person but in the absence of data, we simply assigned different estimates for Φ_B_ to each biting class):

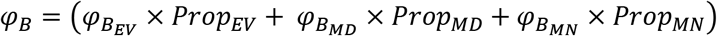

For the outputs in Supplementary Table 6, 𝜑_𝐵_𝐸𝑉__ and 𝜑_𝐵_𝑀𝑁__ were defined as 0.425 (half the contact with bed nets of midnight feeding mosquitoes) whilst 𝜑_𝐵_𝑀𝐷__ was parameterized to be 0.85. This reflected mosquito population that feeds principally in the evening (runs 2, 5, 8, and 11), at midnight (runs 1, 4, 7, and 10) or in the morning (runs 3, 6, 9, and 12).

#### Model simulations

We explored how much biting time might affect estimates of prevalence in 2-10-year-old children in a theoretical high transmission setting i) in the absence of LLINs (runs 1-3 for altered vector competence and runs 7-9 for constant vector competence) and ii) in the presence of LLINs but with equal probability of exposure to LLINs for each of the biting classes (runs 1-3 for altered vector competence and runs 7-9 for constant vector competence); and iii) what would happen if the probability of exposure to LLINs differed between biting classes, consistent with hosts less likely to be in bed and protected by bed nets in the evening and morning (runs 4-6 for altered vector competence and runs 10-12 for constant vector competence). For simplicity, we assumed that: i) the mosquito population is density dependent; ii) biting rates are constant between people; iii) that there are either no interventions, or 50% of people use bed nets, and; iv) there is no pyrethroid resistance in the mosquito population. The scenario was a high transmission, perennial setting such that without interventions prevalence in 2-10-year-old children is about 60%.

We estimated the relative efficacy of LLINs as:

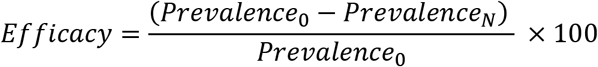

Where subscripts 0 and N represent the scenarios without or with bed nets, respectively. Post bed net prevalence estimates are taken 3 years after LLINs were introduced to estimate the efficacy.

### Statistical analyses

#### Literature review

The ratio of the number of cases where biting time oriented towards either evening or morning (Supplementary Table 1 and Supplementary Table 2) was compared to the expected ratio of 50% using chi-squire goodness-of-fit test. Fisher’s exact test (two-tailed) was used to test if this ratio of evening and morning biting was different between the high and low temperature groups (Supplementary Table 1 and Supplementary Table 2).

#### Mosquito transmission experiments

For analysing infection data in general, Generalized Linear Models (GLM) were used unless otherwise specified. Oocyst intensity data were analysed with a negative binomial error structure with log link considering the highly over-dispersed nature of parasite load data, unless otherwise specified (see Supplementary Table 12). Oocyst or sporozoite prevalence data were analysed with a binomial error structure with logit link. Model fit and distributions were determined based on Akaike’s Information Criterion (AIC) value and residual plots.

For the time-of-day and fluctuating temperature experiment using *A. gambiae,* a Generalized Linear Mixed effects Model (GLMM) was used to examine the effects of time-of-day of blood meal, temperature regime, and their interaction (fixed variables) on oocyst intensity, or oocyst or sporozoite prevalence (dependent variables). Infectious feed was included as a random variable, and dissection day was additionally included as a fixed variable in the model to account for any day effect. Time-of-day groups were pairwise compared for oocyst intensity, or oocyst or sporozoite prevalence within temperature regime groups and between temperature regime groups for each time-of-day group, using post-hoc contrasts followed by Bonferroni corrections.

For the time-of-day and fluctuating temperature experiment using *A. stephensi* replicated in two containers of mosquitoes, GLMM was used to examine the effects of time-of-day of blood meal, temperature regime, and their interaction (fixed variables) on oocyst intensity and prevalence (dependent variables). In addition, in the model analyses, mosquito container and dissection day was included as a random and fixed variable, respectively. For sporozoite prevalence data, GLM was used for pooled data from two containers of mosquitoes after confirming no difference in sporozoite prevalence between two replicate containers using Fisher’s exact test (two-sided) within each treatment group. This was because the variance of the random effect was estimated as zero (i.e. Hessian matrix not positive definite) rendering validity of model uncertain when GLMM was used for prevalence data^84, 85^. Because of slight differences in experimental design, the second time-of-day and fluctuating temperature experiment using *A. stephensi* which used just one container of mosquitoes was analysed separately. Similarly to the model structure used above, GLM was used to examine the effects of time-of-day, temperature regime, and their interaction in addition to dissection day (fixed variables) on oocyst intensity, or oocyst or sporozoite prevalence (dependent variables). For both infection experiments with *A. stephensi*, time-of-day groups were pairwise compared for oocyst intensity, or oocyst or sporozoite prevalence within temperature regime groups and between temperature regime groups for each time-of-day group, using post-hoc contrasts followed by Bonferroni corrections.

For the constant or translocation experiments, GLM was used to examine the effects of temperature treatments, mosquito species, gametocytemia and/or interactions on oocyst intensity, or oocyst or sporozoite prevalence in each study. Treatment groups with zero infections were not included in the analyses^84, 85^. When one control group was compared to all other treatment groups, post-hoc contrasts were used followed by Bonferroni corrections. To examine the effect of parasite intensity on the reduction in infection prevalence at 30°C, a linear regression was used to examine the relationship between the mean oocyst intensity in 27°C control group and per cent reduction in the oocyst prevalence at 30°C. The per cent reduction was calculated as the reduced percentage in oocyst prevalence in the 30°C treatment relative to oocyst prevalence in the 27°C control using the data collected from the infection studies described above.

#### Mosquito translocation and blood feeding compliance

The effects of temperature during blood feeding and time-of-day on the feeding compliance of *A. gambiae* mosquitoes (blood feeding success of individual mosquitoes) were examined by using a GLMM (binomial error structure, logit link) as each treatment group had two technically replicated containers of mosquitoes. Temperature, time-of-day, and their interaction were included as fixed variables, in addition to container of mosquitoes as a random variable in the model.

#### Mosquito translocation and blood meal size

To examine the effect of transferring *A. gambiae* and *A. stephensi* mosquitoes to 27°C for blood feeding from prevailing temperatures at different times of day in 27°C DTR 10°C, GLMM was used for time-of-day, mosquito species, and their interaction as fixed variable, and container of mosquitoes as random variable with normal distribution with an identity link after confirming normality assumptions (e.g. normal distribution of residuals, equal variance, etc.).

#### Thermal avoidance assay

The escape probability of mosquitoes combined from six replicates was analysed using Kaplan Meier Log-rank test to examine the effects of parasite infection on the proportion of mosquitoes that escaped over time. Any mosquitoes that escaped within one minute after opening the gate were left-censored as it was considered a response to human disturbance. Mosquitoes that remained in the holding tube until the end of assay were right-censored.

SPSS Statistics 25 (SPSS Incorporation, Chicago, IL) was used for all analyses. Information on experimental designs, dissection methods, and/or statistical analyses on empirical studies are summarized in Supplementary Table 12.

## Ethical statement

We have compiled with all relevant ethical regulations, and all experiments were conducted under Penn State IBC protocol #48219.

## Reporting summary

Further information on research design is available in the Nature Research Reporting Summary linked to this article.

## Data availability

The authors declare that all data supporting the findings of this study are available within the paper and its supplementary information files.

## Code availability

All code used in modelling analysis is available upon request

## Acknowledgements

We thank Deonna C. Soergel, Janet L. Teeple, and Fhallon Ware-Gilmore for technical assistance, and David A. Kennedy, Eleanore D. Sternberg and Lizhao Ge for advice on statistical analyses. This study was supported by NIH NIAID grant # R01AI110793 and National Science Foundation Ecology and Evolution of Infectious Diseases grant (DEB-1518681). The funders had no role in study design, data collection and analysis, decision to publish, or preparation of the manuscript.

## Author contributions

E.S., J.L.W., E.S.S., T.S.C., and M.B.T. designed research; E.S., J.L.W., N.L.D. and E.S.S. performed research; E.S., M.K.G., E.S.S., and T.S.C. analysed data; and E.S., E.S.S., T.S.C., and M.B.T. wrote the manuscript with inputs from M.K.G., J.L.W, and N.L.D.

