## Supplementary material for "The influence of feeding behaviour and temperature on the capacity of mosquitoes to transmit malaria": SI file

Eunho Suh

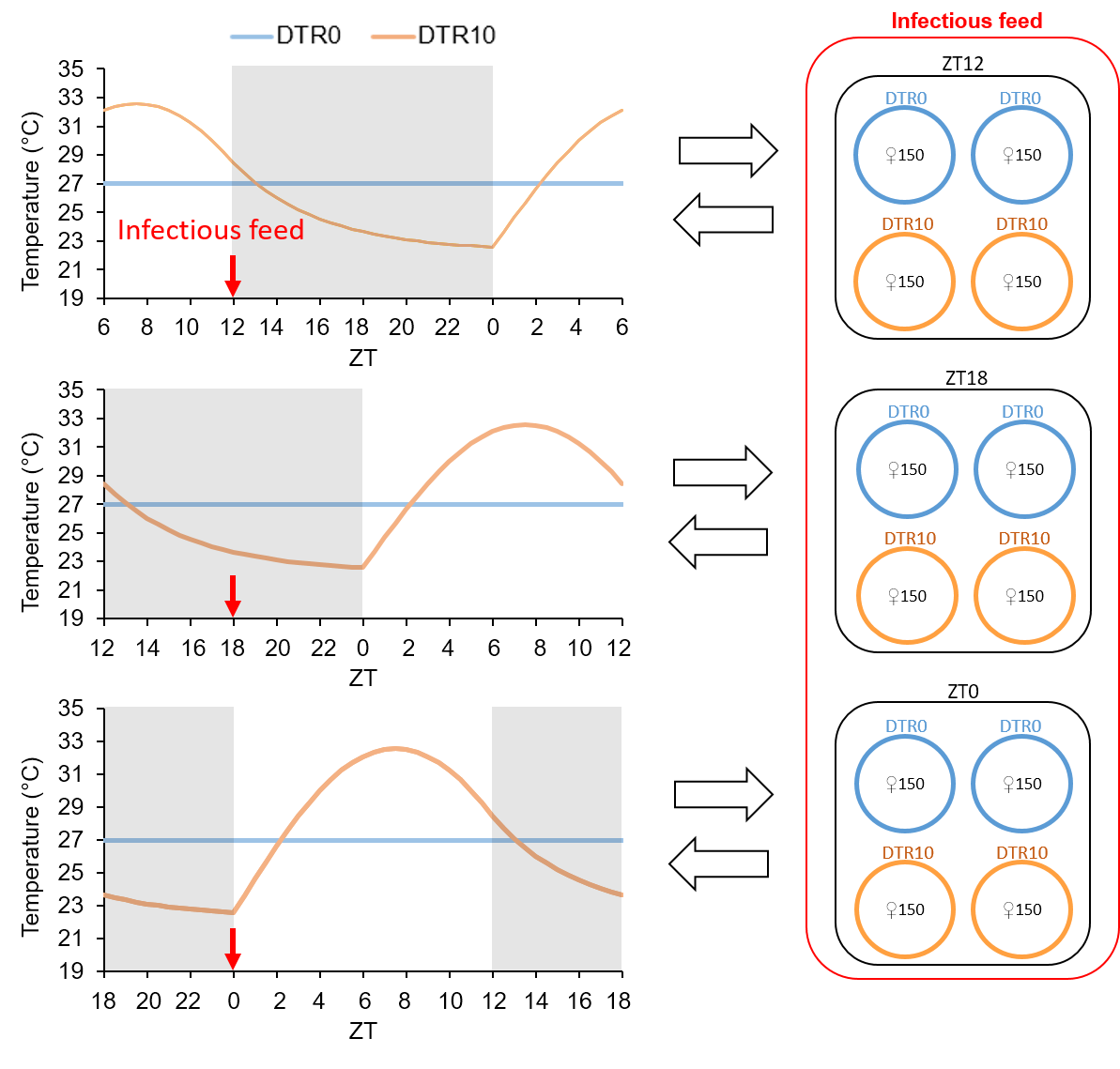

**Supplementary** **Figure 1.** Experimental design for infectious feeds. Adult mosquitoes were acclimated in separate incubators set at either constant (i.e. 27°C with a Diurnal Temperature Range [DTR] of 0°C) or fluctuating (i.e. 27°C with a DTR of 10°C) temperature regimes with a timer offset for each time-of-day treatment so that infectious blood feeding took place simultaneously using the same parasite infected blood meals, but the mosquitoes themselves were at different points in their diel cycle (18:00h [ZT12], 00:00h [ZT18], or 06:00h [ZT0]). Feeding took place in an environmental chamber set at 27°C and then blood fed mosquitoes were immediately moved back to their respective incubators. Each treatment group had 300 female mosquitoes in two containers (150 each) unless otherwise specified.

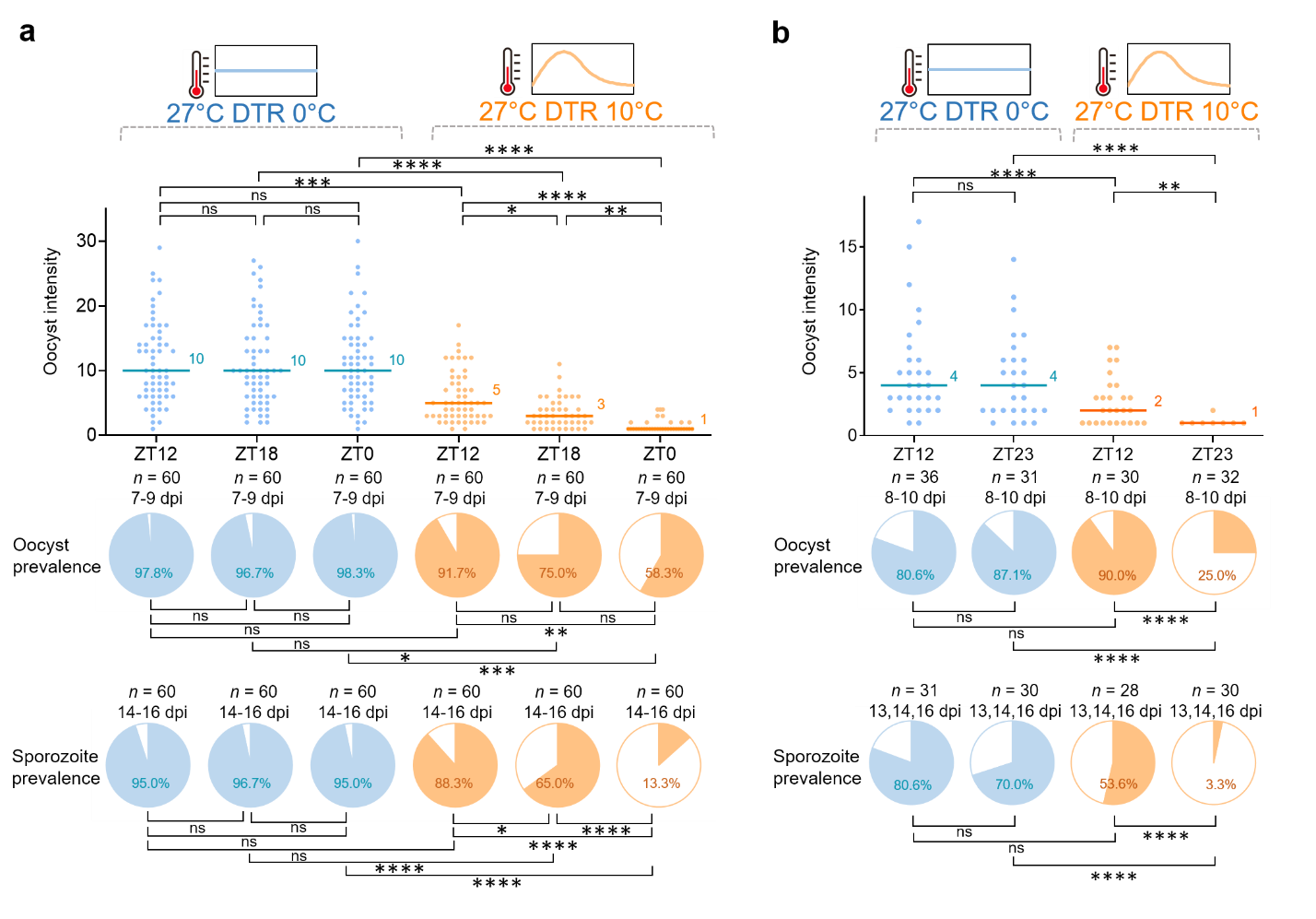

**Supplementary** **Figure 2.** Effects of time-of-day of blood meal and fluctuating temperature on vector competence of *A. stephensi* infected with *P. falciparum* in two separate infection experiments*.* **a,** Mosquitoes were offered infected blood meals at a different time-of-day (18:00h [ZT12], 00:00h [ZT18], or 06:00h [ZT0]) and kept under either constant (i.e. 27°C with a Diurnal Temperature Range [DTR] of 0°C) or fluctuating (i.e. 27°C with a DTR of 10°C) temperature regimes. There is no effect of time-of-day of blood feeding under constant temperature regime (i.e. 27°C DTR 0°C) but vector competence (e.g. sporozoite prevalence) is significantly increased for 18:00h (ZT12) or reduced for 06:00h (ZT0) relative to 00:00h (ZT18) under fluctuating temperature regime (i.e. 27°C DTR 10°C). Results of model analyses to examine the effects of time-of-day and temperature regime on oocyst intensity, or oocyst or sporozoite prevalence are reported in Supplementary Table 4. Twenty mosquitoes were sampled daily for dissecting midguts on 7-9 days post infection (dpi) or salivary glands on 14-16 dpi from two replicate containers (i.e. 10 per each). **b**, Simplified version of time-of-day and fluctuating temperature experiment. There is no effect of time-of-day of blood feeding under constant temperature regime (i.e. 27°C DTR 0°C) but vector competence (e.g. sporozoite prevalence) is significantly reduced for 06:00h (ZT0) under fluctuating temperature regime (i.e. 27°C DTR 10°C). Results of model analyses to examine the effects of time-of-day and temperature regime on oocyst intensity, or oocyst or sporozoite prevalence are reported in Supplementary Table 5. Approximately 10 mosquitoes were sampled daily for dissecting midguts on 8-10 days post infection (dpi) or salivary glands on 13, 14, and 16 dpi. For both (**a)** and (**b),** the scatter plots show oocyst intensity, with the data points representing the number of oocysts found in individual mosquitoes, and the horizontal lines the median. The pie charts show oocyst or sporozoite prevalence calculated as the proportion of infected mosquitoes revealed by dissection of midguts and salivary glands, respectively. *n* indicates the number of mosquito sample per treatment group. Asterisks represent statistically significant difference (* *P* < 0.05, ** *P* < 0.01, *** *P* < 0.001, **** *P* < 0.0001; *P*-values were Bonferroni corrected after pairwise comparisons).

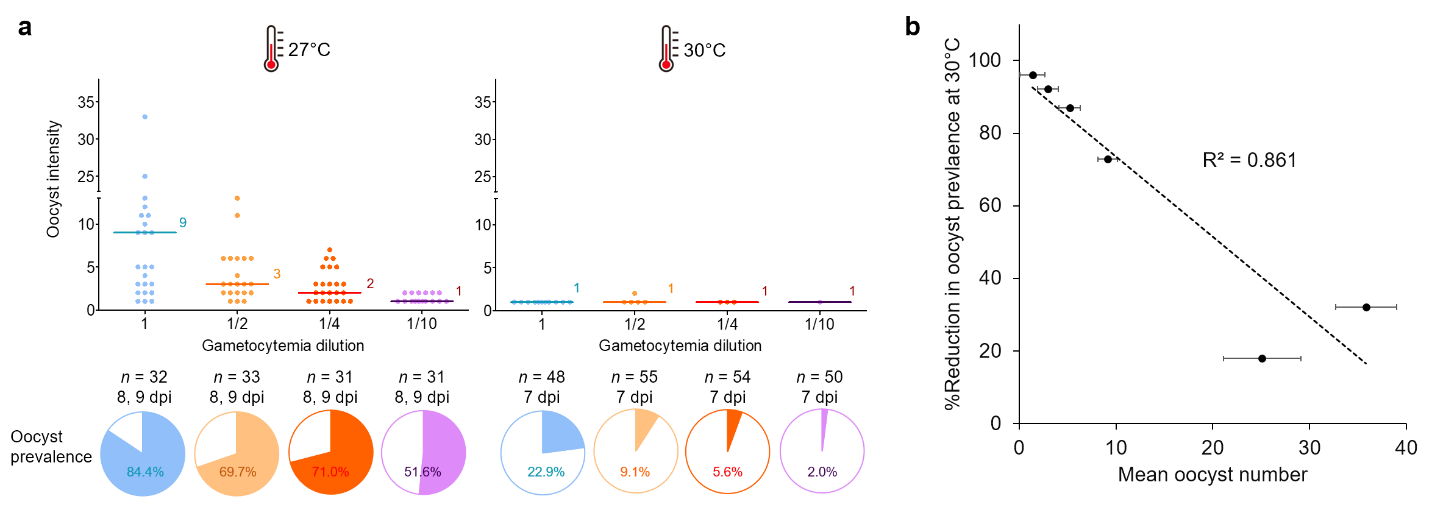

**Supplementary** **Figure 3.** Effects of gametocytemia and temperature on vector competence of *A. stephensi* mosquitoes infected with *P. falciparum* malaria. **a,** Mosquitoes were fed on blood meals with serially diluted gametocytemia (1, 1/2, 1/4, or 1/10) and kept at 27°C or 30°C to examine the effects of high temperature interacting with gametocytemia on oocyst infections. Incubation at 30°C reduces oocyst intensity and prevalence across the board, while oocyst intensity and prevalence is also influenced by gametocytemia. Results of model analyses to examine the effects of gametocytemia and temperature treatment on oocyst intensity or oocyst prevalence are reported in Supplementary Table 8. The scatter plots show oocyst intensity, with the data points representing the number of oocysts found in individual mosquitoes, and the horizontal lines the median. The pie charts show oocyst or sporozoite prevalence calculated as the proportion of infected mosquitoes revealed by dissection of midguts and salivary glands, respectively. *n* indicates the number of mosquito sample per treatment group (dpi = days post infection). **b,** Relationship between per cent reduction in oocyst prevalence due to exposure to 30°C and mean oocyst intensity (error bars = SEM). Per cent reduction represents reduced percentage in oocyst prevalence in the 30°C treatment relative to oocyst prevalence in the 27°C control. Oocyst prevalence and intensity data were derived from experiments reported in Fig. 3a and Supplementary Fig. 3a. The impact of temperature declines as intensity of infection increases. Dashed line indicates linear regression line (*F_1, 4_* = 24.78, *P* = 0.008)

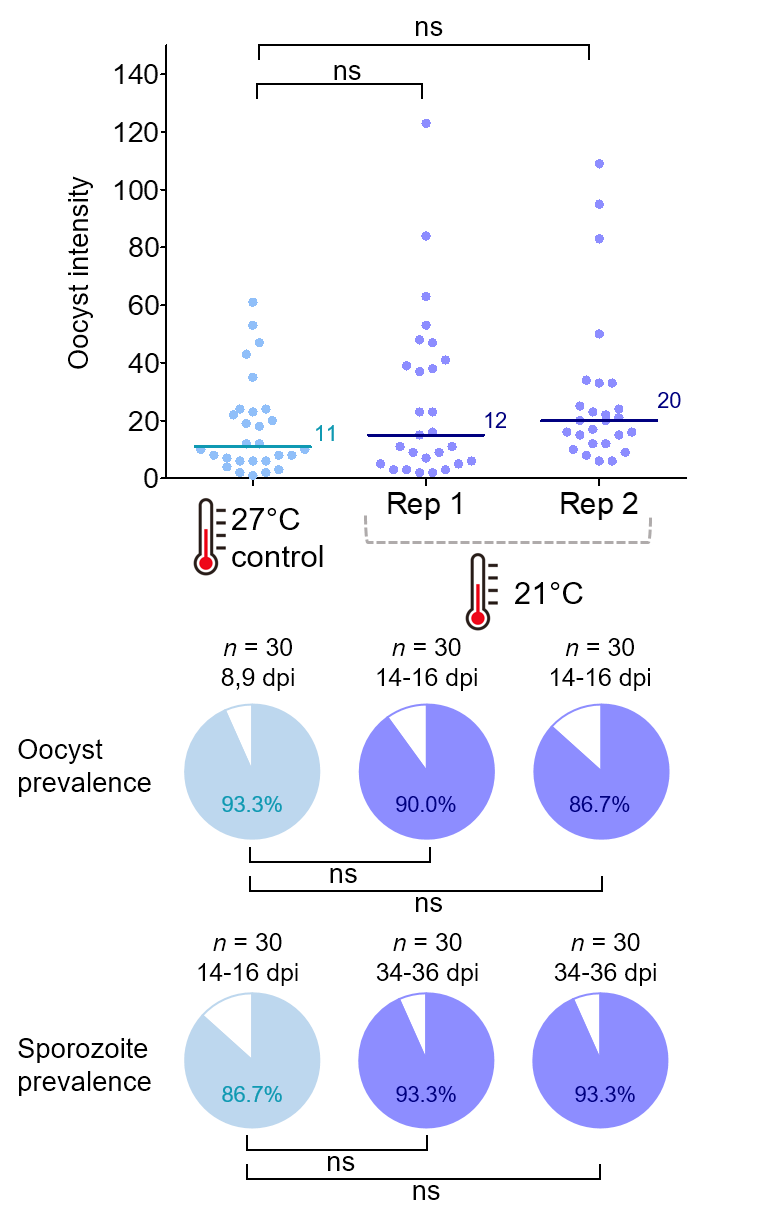

**Supplementary** **Figure 4.** Effect of transferring mosquitoes between 21°C and 27°C on vector competence of *A. gambiae* mosquitoes infected with *P. falciparum* malaria. Treatment mosquitoes in two replicate containers were kept at 21°C, blood fed at 27°C, and moved back to 21°C, while control mosquitoes were kept at 27°C throughout and blood fed at 27°C. Transferring mosquitoes between two different temperatures for blood feeding does not affect vector competence. GLM was used to compare control to each replicate container of mosquitoes with pairwise post-hoc contrasts followed by Bonferroni corrections (ns, not significant at *P* = 0.05). The scatter plots show oocyst intensity, with the data points representing the number of oocysts found in individual mosquitoes, and the horizontal lines the median. The pie charts show oocyst or sporozoite prevalence calculated as the proportion of infected mosquitoes revealed by dissection of midguts and salivary glands, respectively. *n* indicates the number of mosquito sample per treatment (dpi = days post infection).

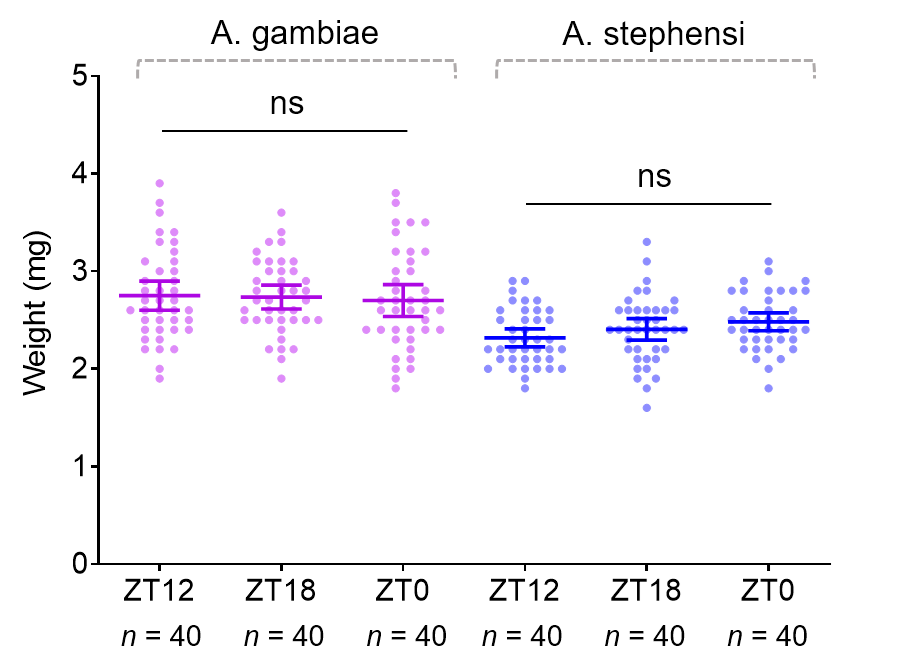

**Supplementary** **Figure 5.** Effects of blood feeding mosquitoes at 27°C transferring from three times-of-day treatments under fluctuating temperature regime (27°C with a DTR of 10°C) on blood meal size of *A. gambiae* and *A. stephensi* mosquitoes. Mosquitoes kept under fluctuating temperature regimes (27°C with a DTR of 10°C) were transferred to 27°C, and offered uninfected blood meals at a different time-of-day (18:00h [ZT12], 00:00h [ZT18], or 06:00h [ZT0]). The whole body weight of blood fed mosquitoes were measured as a proxy for blood meal size. Transferring mosquitoes to 27°C from the prevailing temperature of each time-of-day does not affect blood meal size of mosquitoes. Results of model analyses to examine the effects of species and time-of-day of blood feeding on the body weight are reported in Supplementary Table 11. The scatter plots show body weight of blood fed female mosquitoes. Error bars indicate mean weight with 95% confidence intervals.

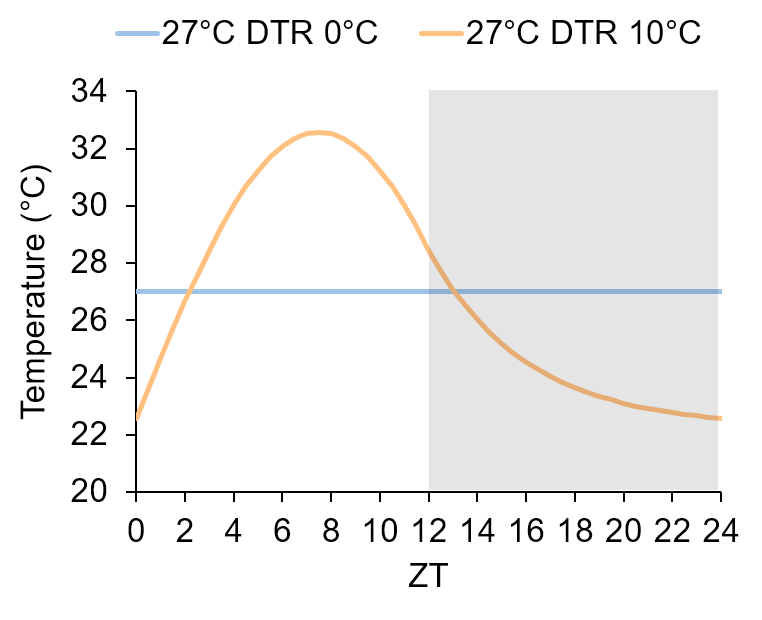

**Supplementary** **Figure 6.** Plots of temperature treatments used in the current study showing 27°C with a Diurnal Temperature Range (DTR) of 0°C or 10°C. The Parton-Logan model was used for the diurnal fluctuating temperature regime that follows a sinusoidal progression and an exponential decay for the day and night cycle, respectively. Shaded areas indicate scotophase.

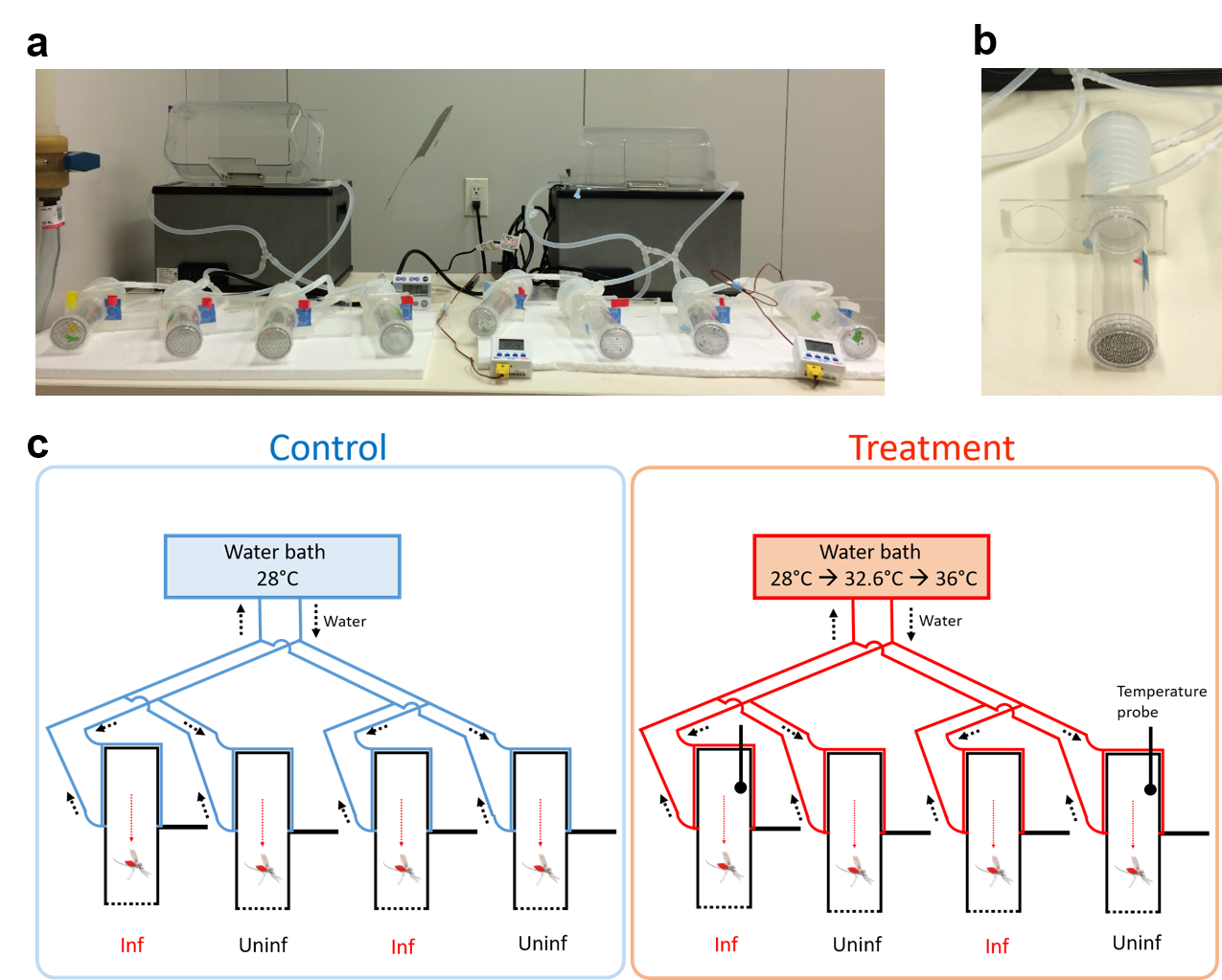

**Supplementary** **Figure 7.** Experimental setup for thermal avoidance assay. Pictures of (**a**) water linked to multiple tubes and (**b**) individual assay tubes. **c**, A schematic diagram of the experimental setup. Total eight assay tubes were used (four for control and four for treatment group) in an assay run, with a total three rounds of assay. Mosquitoes fed with parasite infected (Inf) or uninfected (Uninf) blood meals were introduced into tubes, and the treatments were rotated between the assay rounds.

**Supplementary** **Table 1.** Biting activity profile for *Anopheles* mosquitoes identified to exhibit evening, midnight, or morning biting time in 42 published studies. Biting activities were categorized into ‘evening’, ‘midnight’, or ‘morning’ biting group with peak biting observed before 22:00h, between 22:00 and 05:00h, or after 05:00h, respectively. Studies (i.e. papers reviewed) were grouped into high or low temperature environment (divided by double line in the table).

| Year^ref.^ | Country | Mosquito species | Description on biting activity^a^ | Peak biting time | Temperature (°C)^b^ |
| --- | --- | --- | --- | --- | --- |
| 2017^1^ | Cameroon | *A. gambiae* s.l. | Peak between 00:00 - 02:00 (In+Out), Garoua | Midnight | 29.5 |
|  |  |  | Peak between 22:00 - 00:00 (In+Out), Mayo Oulo | Midnight |  |
|  |  |  | Peak between 00:00 - 02:00 (In+Out), Pitoa | Midnight |  |
|  |  | *A. rufipes* | Peak between 20:00 - 22:00 (In+Out), Garoua | Evening |  |
|  |  |  | Peak between 00:00 - 02:00 (In+Out), Mayo Oulo | Midnight |  |
|  |  |  | Peak between 00:00 - 02:00 (In+Out), Pitoa | Midnight |  |
| 2012^2^ | Benin | *A. funestus* | Peak between 05:00 - 06:00 (In+Out), Lokohoue, 2011 | Morning | 28.7 |
| 2009^3^ | Chad | *A. arabiensis* | Peak between 01:00 - 02:00 (In+Out) | Midnight | 27.9 |
|  |  | *An. pharoensis* | Peak between 21:00 - 22:00 (In+Out) | Evening |  |
|  |  | *A. funestus* | Peak between 03:00 - 04:00 (In+Out) | Midnight |  |
|  |  | *A. ziemanni* | Peak between 18:00 - 19:00 (In+Out) | Evening |  |
| 2017^4^ | Indonesia | *A. vagus* | Peak between 21:00 - 22:00 (In+Out) | Evening | 27.8 |
|  |  | *A. sundaicus* | Peak between 21:00 - 22:00 (In+Out) | Evening |  |
|  |  | *A. subpictus* | Peak between 23:00 - 00:00 (In+Out) | Midnight |  |
|  |  | *A. indefnitus* | Peak between 23:00 - 00:00 (In+Out) | Midnight |  |
|  |  | *A. peditaeniatus* | Peak between 00:00 - 01:00 (In+Out) | Midnight |  |
|  |  | *A. nigerrimus* | Peak between 20:00 - 21:00 (In+Out) | Evening |  |
| 2011^5^ | Solomon Islands | *A. farauti* | Peak between 18:00 - 19:00 (In), Pala, Dec 2010 | Evening | 27.1 |
|  |  |  | Peak between 19:00 - 21:00 (Out), Pala, Dec 2010 | Evening |  |
| 2007^6^ | Tanzania | *A. gambiae* s.s. | Peak between 01:00 - 02:00 (In) | Midnight | 27.0 |
|  |  | *A. gambiae* s.s. | Peak between 01:00 - 02:00 (Out) | Midnight |  |
|  |  | *A. arabiensis* | Peak between 20:00 - 21:00 (In) | Evening |  |
|  |  | *A. arabiensis* | Peak between 22:00 - 23:00 (Out) | Midnight |  |
| 2008^7^ | French Guiana | *A. darlingi* | Peak between 22:30 - 23:30 (Out), Twenké | Midnight | 27.0 |
|  |  |  | Peak between 22:30 - 23:30 (Out), Taluéne | Midnight |  |
|  |  |  | Peak between 05:30 - 06:30 (Out), Cayodé | Morning |  |
| 2012^8^ | Suriname | *A. darlingi* | Peak between 05:00 - 06:00 (In), Drietabiki | Morning | 27.0 |
|  |  |  | Peak between 04:00 - 05:00 (Out), Drietabiki | Midnight |  |
|  |  |  | Peak between 01:00 - 02:00 (In), Jamaica | Midnight |  |
|  |  |  | Peak between 01:00 - 02:00 (Out), Jamaica | Midnight |  |
| 2014^9^ | Senegal | *A. funestus* | Peak between 08:00 - 09:00 (In+Out) | Morning | 27.0 |
| 2004^10^ | Eritrea | *A. gambiae* s.l. | Peak between 02:00 - 03:00 (In), Gash-barka | Midnight | 26.8 |
|  |  |  | Peak between 22:00 - 23:00 (Out), Gash-barka | Midnight |  |
|  |  |  | Peak between 02:00 - 03:00 (In), Debub | Midnight |  |
|  |  |  | Peak between 21:00 - 22:00 (Out), Debub | Evening |  |
|  |  |  | Peak between 01:00 - 02:00 (In), Anseba | Midnight |  |
|  |  |  | Peak between 21:00 - 22:00 (Out), Anseba | Evening |  |
| 2012^11^ | Cameroon | *A. gabmiae* s.l. | Peak between 01:00 - 02:00 (Out) | Midnight | 26.8 |
| 2017^12^ | Papua New Guinea | *A. farauti* 4 | peak between 19:00 - 20:00 (Out), Kokofine, 2011 | Evening | 26.7 |
|  |  |  | peak between 20:00 - 21:00 (Out), Mauno, 2011 | Evening |  |
| 2005^13^ | Bolivia | *A. darlingi* | Peak between 20:00 - 21:00 (Out) | Evening | 26.6 |
| 2011^14^ | Solomon Islands | *A. farauti* | Peak between 18:00 - 19:00 (In) | Evening | 26.5 |
|  |  |  | Peak between 19:00 - 20:00 (Out) | Evening |  |
|  |  | *A. solomonis* | Peak between 18:00 - 19:00 (In) | Evening |  |
|  |  |  | Peak between 18:00 - 19:00 (Out) | Evening |  |
| 2017^15^ | Tanzania | *An. arabiensis* | Peak between 20:00 - 21:00 (Out) | Evening | 26.5 |
|  |  | *An. funestus* | Peak between 05:00 - 06:00 (Out) | Morning |  |
| 2008^16^ | Ghana | *A. gambiae* s.l. | Peak between 02:00 - 03:00 (Out), Dzorwulu | Midnight | 26.4 |
|  |  |  | Peak between 03:00 - 04:00 (Out), Kaneshie | Midnight |  |
|  |  |  | Peak between 02:00 - 03:00 (Out), Korle Bu | Midnight |  |
|  |  |  | Peak between 02:00 - 03:00 (Out), Kotobabi | Midnight |  |
|  |  |  | Peak between 02:00 - 03:00 (Out), La | Midnight |  |
|  |  |  | Peak between 03:00 - 04:00 (Out), Ushertown | Midnight |  |
| 2013^17^ | Uganda | *A. gambiae* s.l. | Peak between 04:00 - 05:00 (In+Out), Bugabula | Midnight | 26.2 |
|  |  |  | Peak between 02:00 - 03:00 (In+Out), Budiope | Midnight |  |
|  |  | *A. funestus* | Peak between 04:00 - 05:00 (In+Out), Budiope | Midnight |  |
|  |  |  | Peak between 05:00 - 06:00 (In+Out), Bugabula | Morning |  |
| 2015^18^ | Peru | *A. darlingi* | Peak between 21:00 - 22:00 (Out), Riverine | Evening | 26.1 |
|  |  |  | Peak between 22:00 - 23:00 (Out), Highway | Midnight |  |
| 2015^19^ | Peru | *A. darlingi* | Peak between 18:00 - 19:00 (Out), San José de Lupuna, April 2011 | Evening | 26.1 |
|  |  |  | Peak between 23:00 - 00:00 (Out), Villa del Buen Pastor, April 2011 | Midnight |  |
|  |  |  | Peak between 22:00 - 23:00 (Out), Cahuide, May 2012 | Midnight |  |
| 2009^20^ | Colombia | *A. darlingi* | Peak between 18:00 - 19:00 (In) | Evening | 26.0 |
|  |  |  | Peak between 20:00 - 21:00 (Out) | Evening |  |
|  |  | *A. oswaldoi* | Peak between 18:00 - 19:00 (Out) | Evening |  |
|  |  |  | Peak between 18:00 - 19:00 (Out) | Evening |  |
| 2014^21^ | Solomon Islands | *A. farauti* | Peak between 19:00 - 20:00 (In) | Evening | 26.0 |
|  |  |  | Peak between 19:00 - 20:00 (Out) | Evening |  |
| 2015^22^ | Equatorial Guinea | *Anopheles* spp. | Peak between 03:00 - 04:00 (In) | Midnight | 26.0 |
|  |  |  | Peak between 03:00 - 04:00 (Out) | Midnight |  |
| 2016^23^ | Solomon Islands | *A. farauti* s.s. | Peak between 18:00 - 19:00 (In) | Evening | 26.0 |
|  |  |  | Peak between 18:00 - 19:00 (Out) | Evening |  |
| 2012^24^ | Zambia | *A. funestus* | Peak between 03:00 - 04:00 (In), LLINs alone | Midnight | 25.7 |
|  |  |  | Peak between 05:00 - 06:00 (Out), LLINs alone | Morning |  |
|  |  |  | Peak between 03:00 - 04:00 (In), LLINs + IRS | Midnight |  |
|  |  |  | Peak between 00:00 - 01:00 (In), LLINs + IRS | Midnight |  |
|  |  | *A. quadriannulatus* | Peak between 19:00 - 20:00 (In), LLINs alone | Evening |  |
|  |  |  | Peak between 20:00 - 21:00 (Out), LLINs alone | Evening |  |
|  |  |  | Peak between 20:00 - 21:00 (In), LLINs + IRS | Evening |  |
|  |  |  | Peak between 21:00 - 22:00 (Out), LLINs + IRS | Evening |  |
| 2011^25^ | Equatorial Guinea | *A. gambiae* s.s. | Peak between 23:00 - 00:00 (In) | Midnight | 24.6 |
|  |  |  | Peak between 23:00 - 00:00 (Out) | Midnight |  |
| 2011^26^ | Indonesia | *A. aconitus* | Peak between 22:00 - 23:00 (In+Out) | Midnight | 24.1 |
|  |  | *A. vagus* | Peak between 19:00 - 20:00 (In+Out), West Timor | Evening |  |
|  |  | *A. barbirostris* | Peak between 01:00 - 02:00 (In+Out) | Midnight |  |
|  |  | *A. vagus* | Peak between 02:00 - 03:00 (In+Out), Java | Midnight |  |
|  |  | *A. subpictus* | Peak between 22:00 - 23:00 (In+Out), West Timor | Midnight |  |
| 2007^27^ | Venezuela | *A. darlingi* | Peak between 01:00 - 02:00 (In) | Midnight | 24.0 |
| 2012^28^ | Iran | *A. culcifacies* | Peak between 23:00 - 00:00 (In+Out) | Midnight | 23.5 |
|  |  | *A. fluviatilis* | Peak between 22:00 - 23:00 (In+Out) | Midnight |  |
|  |  | *A. stephensi* | Peak between 19:00 - 20:00 (In+Out) | Evening |  |
| 2011^29^ | Tanzania | *A. gambiae* s.l. | Peak between 00:00 - 01:00 (In), 2009 | Midnight | 23.3 |
|  |  |  | Peak between 22:00 - 23:00 (Out), 2009 | Midnight |  |
|  |  | *A. funestus* | Peak between 20:00 - 21:00 (In), 2009 | Evening |  |
|  |  |  | Peak between 22:00 - 23:00 (Out), 2009 | Midnight |  |
| 2005^30^ | India | *A. baimaii* | Peak between 22:00 - 23:00 (In) | Midnight | 23.2 |
| 2001^31^ | Kenya | *A. gambiae* s.l. | Peak between 23:00 - 00:00 (Out), bed net village | Midnight | 23.0 |
|  |  |  | Peak between 23:00 - 00:00 (Out), control village | Midnight |  |
|  |  | *A. funestus* | Peak between 22:00 - 23:00 (Out), bed net village | Midnight |  |
|  |  |  | Peak between 23:00 - 00:00 (Out), control village | Midnight |  |
| 2000^32^ | Mozambique | *A. arabiensis* | Peak between 01:00 - 02:00 (In) | Midnight | 22.8 |
|  |  |  | Peak between 23:00 - 00:00 (Out) | Midnight |  |
|  |  | *A. funestus* | Peak between 02:00 - 03:00 (In) | Midnight |  |
|  |  |  | Peak between 02:00 - 03:00 (Out) | Midnight |  |
| 2015^33^ | Uganda | *A. gambiae* s.l. | Peak between 19:00 - 20:00 (In+Out), Engari, rainy season | Evening | 22.3 |
|  |  |  | Peak between 19:00 - 20:00 (In+Out), Engari, dry season | Evening |  |
|  |  |  | Peak between 19:00 - 20:00 (In+Out), Kigorogoro, dry season | Evening |  |
| 2015^34^ | Kenya | *A. gambiae* s.l. | Peak between 22:00 - 00:00 (In) | Midnight | 22.1 |
|  |  |  | Peak between 18:00 - 20:00 (Out) | Evening |  |
|  |  | *A. funestus* | Peak between 18:00 - 20:00 (In) | Evening |  |
|  |  |  | Peak between 18:00 - 20:00 (Out) | Evening |  |
| 2006^35^ | Tanzania | *A. gambiae* s.l. | Peak between 05:00 - 06:00 (In), Lupiro 2004 | Morning | 21.8 |
|  |  |  | Peak between 23:00 - 00:00 (Out), Lupiro 2004 | Midnight |  |
| 2014^36^ | Kenya | *A. gambiae* s.s. | Peak between 05:00 - 06:00 (In), Asembo 2011 | Morning | 21.8 |
|  |  |  | Peak between 01:00 - 02:00 (Out), Asembo 2011 | Midnight |  |
|  |  | *A. arabiensis* | Peak between 01:00 - 02:00 (In), Asembo 2011 | Midnight |  |
|  |  |  | Peak between 03:00 - 04:00 (Out), Asembo 2011 | Midnight |  |
|  |  | *A. funestus* | Peak between 01:00 - 02:00 (In), Asembo 2011 | Midnight |  |
|  |  |  | Peak between 01:00 - 02:00 (Out), Asembo 2011 | Midnight |  |
| 2015^37^ | Madagascar | *A. coustani* | Peak between 22:00 - 23:00 (In) | Midnight | 21.3 |
|  |  |  | Peak between 19:00 - 21:00 (Out) | Evening |  |
|  |  | *A. mascarensis* | Peak between 01:00 - 02:00 (In) | Midnight |  |
|  |  |  | Peak between 02:00 - 03:00 (Out) | Midnight |  |
|  |  | *A. funestus* | Peak between 21:00 - 22:00 (In) | Evening |  |
|  |  |  | Peak between 02:00 - 03:00 (Out) | Midnight |  |
|  |  | *A. arabiensis* | Peak between 04:00 - 05:00 (In) | Midnight |  |
|  |  |  | Peak between 00:00 - 01:00 (Out) | Midnight |  |
| 2016^38^ | Ethiopia | *A. arabiensis* | Peak between 00:00 - 01:00 (In) | Midnight | 21.1 |
|  |  |  | Peak between 21:00 - 22:00 (Out) | Evening |  |
|  |  | *A. pharoensis* | Peak between 19:00 - 20:00 (In) | Evening |  |
|  |  |  | Peak between 19:00 - 20:00 (Out) | Evening |  |
|  |  | *A. ziemanni* | Peak between 19:00 - 20:00 (In) | Evening |  |
|  |  |  | Peak between 19:00 - 20:00 (Out) | Evening |  |
|  |  | *A. funestus* s.l. | Peak between 23:00 - 00:00 (In) | Midnight |  |
|  |  |  | Peak between 21:00 - 22:00 (Out) | Evening |  |
| 2010^39^ | Ethiopia | *A. arabiensis* | Peak between 19:00 - 20:00 (In) | Evening | 20.0 |
|  |  |  | Peak between 18:00 - 19:00 (Out) | Evening |  |
|  |  | *A. pharoensis* | Peak between 20:00 - 21:00 (In) | Evening |  |
|  |  |  | Peak between 19:00 - 20:00 (Out) | Evening |  |
|  |  | *A. coustani* | Peak between 18:00 - 19:00 (In) | Evening |  |
|  |  |  | Peak between 18:00 - 19:00 (Out) | Evening |  |
| 2010^40^ | Zambia | *A. arabiensis* | Peak between 24:00 - 01:00 (In) | Midnight | 19.9 |
|  |  |  | Peak between 01:00 - 02:00 (Out) | Midnight |  |
| 2016^41^ | Ethiopia | *A. gambiae* s.l. | Peak between 19:00 - 20:00 (In) | Evening | 18.0 |
|  |  |  | Peak between 19:00 - 20:00 (Out) | Evening |  |
|  |  | *A. coustani* s.l. | Peak between 21:00 - 22:00 (In) | Evening |  |
|  |  |  | Peak between 19:00 - 20:00 (Out) | Evening |  |
|  |  | *A. pharoensis* | Peak between 19:00 - 20:00 (In) | Evening |  |
|  |  |  | Peak between 20:00 - 21:00 (Out) | Evening |  |
| 2012^42^ | Ethiopia | *A. arabiensis* | Peak between 19:00 - 20:00 (In) | Evening | 17.4 |
|  |  |  | Peak between 19:00 - 20:00 (Out) | Evening |  |

In: peak biting observed for indoor biting.

Out: peak biting observed for outdoor biting.

In+Out: peak biting observed for combined data of indoor and outdoor biting.

^a^If a subset of data showed a shift in biting time in each study, the data set was described for the details such as study sites, year, and/or intervention methods (e.g., long-lasting insecticide-treated bed nets [LLINs], indoor residual spray [IRS], etc.).

^b^Temperature measures represent monthly mean temperature of regional estimates for the study sites and study periods in each paper reviewed, otherwise specified in each paper.

**Supplementary** **Table 2.** Summary of biting activity profile from Supplementary Table 1

| Biting time | No. cases^a^ (%) by temperature measured^b^ | |
| --- | --- | --- |
|  | High (25°C or above) | Low (< 25°C) |
| Evening | 33 (21.9) | 31 (20.5) |
| Midnight | 40 (26.5) | 38 (25.2) |
| Morning | 7 (4.6) | 2 (1.3) |

^a^A ‘case’ indicates a mosquito species or species group determined for biting activity in Supplementary Table 1.

^b^Temperature measured indicates the representative temperature data for each study reviewed in Supplementary Table 1.

**Supplementary** **Tables 3.** GLMMs examining the effects of time-of-day (18:00h [ZT12], 00:00h [ZT18], and 06:00h [ZT0]) and temperature regime (27°C DTR 0°C and 27°C DTR 10°C) on oocyst intensity, or oocyst or sporozoite prevalence in *A. gambiae* (See Fig. 1)

|  | | Oocyst intensity | | Oocyst prevalence | | Sporozoite prevalence | |
| --- | --- | --- | --- | --- | --- | --- | --- |
| Effect | *df* | *F* | *P* | *F* | *P* | *F* | *P* |
| Time^a^ | 2 | 9.91 | < 0.0001 | 13.42 | < 0.0001 | 17.48 | < 0.0001 |
| DTR^b^ | 1 | 93.02 | < 0.0001 | 74.63 | < 0.0001 | 47.96 | < 0.0001 |
| Time × DTR | 2 | 17.36 | < 0.0001 | 18.64 | < 0.0001 | 16.19 | < 0.0001 |
| Day^c^ | 2 | 0.83 | 0.436 | 0.06 | 0.940 | 2.51 | 0.088 |

*LR*-*χ^2^*: Likelihood ratio chi-square value.

^a^Time-of-day.

^b^Diurnal temperature range.

^c^Dissection day (day post infection).

**Supplementary** **Tables 4.** Model analyses examining the effects of time-of-day (18:00h [ZT12], 00:00h [ZT18], and 06:00h [ZT0]) and temperature regime (27°C DTR 0°C and 27°C DTR 10°C) on oocyst intensity (GLMM), or oocyst (GLMM) or sporozoite prevalence (GLM) in *A. stephensi* (see Supplementary Fig. 2a)

|  | | Oocyst intensity | | Oocyst prevalence | | Sporozoite prevalence | |
| --- | --- | --- | --- | --- | --- | --- | --- |
| Effect | *df* | *F* | *P* | *F* | *P* | *LR*-*χ^2^* | *P* |
| Time^a^ | 2 | 15.07 | < 0.0001 | 1.20 | 0.318 | 13.00 | 0.002 |
| DTR^b^ | 1 | 158.25 | < 0.0001 | 18.95 | < 0.001 | 59.59 | < 0.0001 |
| Time × DTR | 2 | 13.23 | < 0.0001 | 0.97 | 0.393 | 14.08 | < 0.001 |
| Day^c^ | 2 | 0.21 | 0.812 | 0.12 | 0.890 | 1.79 | 0.410 |

*LR*-*χ^2^*: Likelihood ratio chi-square value.

^a^Time-of-day.

^b^Diurnal temperature range.

^c^Dissection day (day post infection).

**Supplementary** **Table 5.** GLMs examining the effects of time-of-day (18:00h [ZT12] and 05:00h [ZT23]) and temperature regime (27°C DTR 0°C and 27°C DTR 10°C) on oocyst intensity, or oocyst or sporozoite prevalence in *A. stephensi* (see Supplementary Fig. 2b)

|  | | Oocyst intensity | | Oocyst prevalence | | Sporozoite prevalence | |
| --- | --- | --- | --- | --- | --- | --- | --- |
| Effect | *df* | *LR*-*χ^2^* | *P* | *LR*-*χ^2^* | *P* | *LR*-*χ^2^* | *P* |
| Time^a^ | 1 | 9.31 | 0.002 | 8.17 | 0.004 | 16.01 | < 0.0001 |
| DTR^b^ | 1 | 45.64 | <0.0001 | 4.93 | 0.026 | 33.29 | < 0.0001 |
| Time × DTR | 1 | 4.78 | 0.029 | 16.51 | < 0.0001 | 7.38 | 0.007 |
| Day^c^ | 2 | 10.65 | 0.005 | 2.35 | 0.309 | 0.80 | 0.672 |

*LR*-*χ^2^*: Likelihood ratio chi-square value.

^a^Time-of-day.

^b^Diurnal temperature range.

^c^Dissection day (day post infection).

**Supplementary** **Table 6.** Outputs from a malaria transmission dynamics model illustrating the potential effect of altered or constant vector competence in mosquitoes biting in the evening (EV), at midnight (MD), or in the morning (MN) on malaria prevalence and efficacy of bed nets (LLINs). Post bed net prevalence estimates are taken 3 years after they were introduced at 50% usage and maintained annually to estimate the efficacy of LLINs (See Fig. 2)

| Run | Vector competence | Proportion of mosquitoes biting during different periods of the night | | | Proportion of bites received in bed | | | Prevalence (%) in 2 – 10-year old children^§^ | | Estimated efficacy of LLINs (% relative reduction in prevalence)^§^ |
| --- | --- | --- | --- | --- | --- | --- | --- | --- | --- | --- |
|  |  | EV | MD | MN | EV | MD | MN | Without LLINs | With LLINs |  |
| 1 | Altered^¶^ | 0.15 | 0.7 | 0.15 | 0.85^†^ | 0.85^†^ | 0.85^†^ | 59.5 (54.4 – 63.7) | 15.6 (11.5 – 19.8) | 73.7 (69.0 - 78.9) |
| 2 | Altered^¶^ | 0.7 | 0.3 | 0 | 0.85^†^ | 0.85^†^ | 0.85^†^ | 68.5 (65.6 – 70.8) | 25.2 (21.8 – 28.1) | 63.3 (66.8 – 60.3) |
| 3 | Altered^¶^ | 0 | 0.3 | 0.7 | 0.85^†^ | 0.85^†^ | 0.85^†^ | 39.0 (30.4 – 48.6) | 3.4 (1.4 – 7.6) | 91.3 (84.3 – 95.3) |
| 4 | Altered^¶^ | 0.15 | 0.7 | 0.15 | 0.43 | 0.85^†^ | 0.43 | 59.5 (54.4 – 63.7) | 22.6 (15.6 – 28.1) | 62.0 (56.8 - 67.9) |
| 5 | Altered^¶^ | 0.7 | 0.3 | 0 | 0.43 | 0.85^†^ | 0.43 | 68.5 (65.6 – 70.8) | 44.4 (40.4 – 47.6) | 35.3 (38.4 – 32.8) |
| 6 | Altered^¶^ | 0 | 0.3 | 0.7 | 0.43 | 0.85^†^ | 0.43 | 39.0 (30.4 – 48.6) | 12.1 (6.4 -20.7) | 68.9 (57.4 – 78.9) |
| 7 | Constant^ǂ^ | 0.15 | 0.7 | 0.15 | 0.85^†^ | 0.85^†^ | 0.85^†^ | 58.4 (52.2 – 63.3) | 14.7 (9.9 – 19.3) | 74.9 (69.5 – 81.1) |
| 8 | Constant^ǂ^ | 0.7 | 0.3 | 0 | 0.85^†^ | 0.85^†^ | 0.85^†^ | 58.4 (52.2 – 63.3) | 14.7 (9.9 – 19.3) | 74.9 (69.5 – 81.1) |
| 9 | Constant^ǂ^ | 0 | 0.3 | 0.7 | 0.85^†^ | 0.85^†^ | 0.85^†^ | 58.4 (52.2 – 63.3) | 14.7 (9.9 – 19.3) | 74.9 (81.1 – 69.5) |
| 10 | Constant^ǂ^ | 0.15 | 0.7 | 0.15 | 0.43 | 0.85^†^ | 0.43 | 58.4 (52.2 – 63.3) | 21.4 (15.4 – 27.0) | 63.3 (57.4 – 70.5) |
| 11 | Constant^ǂ^ | 0.7 | 0.3 | 0 | 0.43 | 0.85^†^ | 0.43 | 58.4 (52.2 – 63.3) | 31.4 (24.4 – 37.4) | 46.3 (53.3 – 40.9) |
| 12 | Constant^ǂ^ | 0 | 0.3 | 0.7 | 0.43 | 0.85^†^ | 0.43 | 58.4 (52.2 – 63.3) | 31.4 (24.4 – 37.4) | 46.3 (53.3 – 40.9) |

^¶^Vector competence is assumed to be increased, intermediate, or low for mosquitoes biting in the evening, at midnight, or in the morning, respectively.

^ǂ^Vector competence is assumed to be equal with respect to biting time.

^†^See reference *A. gambiae* s.s.^43^.

^§^Numbers in parentheses represent 95% confidence intervals.

**Supplementary** **Table 7.** GLMs examining the effects of mosquito species (*A. gambiae* and *A. stephensi*) and/or temperature treatment (27°C and 30°C) on oocyst intensity or oocyst prevalence (see Fig. 3a). Oocyst prevalence data were pooled within each temperature treatment group after confirming no difference between two species (Fisher’s exact test, two-sided, *P* > 0.05) to ensure model validity^44^^,^^45^

|  | | Oocyst intensity | | Oocyst prevalence | |
| --- | --- | --- | --- | --- | --- |
| Effect | *df* | *LR*-*χ^2^* | *P* | *LR*-*χ^2^* | *P* |
| Species | 1 | 0.23 | 0.632 | NA | NA |
| Temperature | 1 | 78.7 | < 0.0001 | 36.9 | < 0.0001 |
| Species × Temperature | 1 | 1.29 | 0.256 | NA | NA |

*LR*-*χ^2^*: Likelihood ratio chi-square value.

**Supplementary** **Table 8.** GLMs examining the effects of gametocytemia dilutions (1, 1/2, 1/4, and 1/10) and temperature treatment (27°C and 30°C) on oocyst intensity or oocyst prevalence in *A. stephensi* (see Supplementary Fig. 3a)

|  | | Oocyst intensity | | Oocyst prevalence | |
| --- | --- | --- | --- | --- | --- |
| Effect | *df* | *LR*-*χ^2^* | *P* | *LR*-*χ^2^* | *P* |
| Gametocytemia | 3 | 2.48 | 0.479 | 20.3 | < 0.0001 |
| Temperature | 1 | 5.96 | 0.015 | 138 | < 0.0001 |
| Gametocytemia × Temperature | 1 | 2.72 | 0.438 | 1.33 | 0.724 |

*LR*-*χ^2^*: Likelihood ratio chi-square value.

**Supplementary** **Table 9.** Blood feeding compliance of *A. gambiae* mosquitoes fed at either 27°C or 21°C. Mosquitoes were kept at either 27°C DTR 0°C or 21°C DTR 0°C and fed infectious blood meals at a different time-of-day (18:00h [ZT12], 00:00h [ZT18], or 06:00h [ZT0]) at their corresponding temperature (i.e. either 27°C or 21°C). Data for feeding compliance at 27°C were obtained from the infectious feed (2^nd^ feed) reported in Fig. 1 (i.e. 27°C DTR 0°C treatment group), and data for feeding compliance at 21°C were obtained from a separate infectious feed. GLMM examining the effects of temperature and time-of-day on the feeding compliance is reported in Supplementary Table 10

| Blood feeding temperature | Time-of-day | No. fed | No. total | % fed |
| --- | --- | --- | --- | --- |
| 27°C | ZT12 | 117 | 118 | 99.2 |
|  |  | 116 | 118 | 98.3 |
|  | ZT18 | 113 | 115 | 98.3 |
|  |  | 114 | 116 | 98.3 |
|  | ZT0 | 112 | 116 | 96.6 |
|  |  | 112 | 117 | 95.7 |
| 21°C | ZT12 | 112 | 119 | 94.1 |
|  |  | 108 | 118 | 91.5 |
|  | ZT18 | 115 | 118 | 97.5 |
|  |  | 108 | 117 | 92.3 |
|  | ZT0 | 115 | 115 | 100.0 |
|  |  | 113 | 117 | 96.6 |

**Supplementary** **Table 10.** GLMM examining the effects of blood feeding temperature (27°C and 21°C) and time-of-day (18:00h [ZT12], 00:00h [ZT18], and 06:00h [ZT0]) on feeding compliance (See Supplementary Table 9)

|  | | Feeding compliance | |
| --- | --- | --- | --- |
| Effect | *df* | *F* | *P* |
| Temperature | 1 | 3.05 | 0.131 |
| Time-of-day | 2 | 0.08 | 0.926 |
| Temperature × Time-of-day | 2 | 3.98 | 0.080 |

**Supplementary** **Table 11.** GLMM examining the effects of species (*A. gambiae* and *A. stephensi*) and time-of-day (18:00h [ZT12], 00:00h [ZT18], and 06:00h [ZT0]) on body weight of blood fed mosquitoes (See Supplementary Fig. 5)

|  | | Feeding compliance | |
| --- | --- | --- | --- |
| Effect | *F* | *F* | *P* |
| Species | 1 | 43.09 | < 0.0001 |
| Time-of-day | 2 | 0.46 | 0.635 |
| Species × Time-of-day | 2 | 1.56 | 0.213 |

**Supplementary Table 12.** Summary of experiment design, dissection method, and/or statistical model analysis for empirical studies (additional information are available in the main text)

| Reference (description on experiment) | Mosquito dissection | Treatment | | Sample size | Dpi^†^ | # Replicate container | # Mosquito per container | Model analysis | Dependent variables | Model structure and explanatory variables | Error structure and link for dependent variables |
| --- | --- | --- | --- | --- | --- | --- | --- | --- | --- | --- | --- |
| Fig. 1 and Supplementary Table 3 (effects of time-of-day and fluctuating temperature on vector competence in *A. gambiae*) | Midguts | 27°C DTR 0°C | ZT12 | 120 | 7-9 | 4 | 150 or 120^ǂ^ | GLMM | Oocyst intensity, or oocyst or sporozoite prevalence | Time-of-day + Temperature regime + Time-of-day × Temperature regime + Dissection day + Infectious feed^*^ | Oocyst intensity - negative binomial distribution with log link; Oocyst and sporozoite prevalence - binomial distribution with logit link |
|  |  |  | ZT18 | 120 | 7-9 | 4 | 150 or 120^ǂ^ |  |  |  |  |
|  |  |  | ZT0 | 120 | 7-9 | 4 | 150 or 120^ǂ^ |  |  |  |  |
|  |  | 27°C DTR 10°C | ZT12 | 120 | 7-9 | 4 | 150 or 120^ǂ^ |  |  |  |  |
|  |  |  | ZT18 | 120 | 7-9 | 4 | 150 or 120^ǂ^ |  |  |  |  |
|  |  |  | ZT0 | 120 | 7-9 | 4 | 150 or 120^ǂ^ |  |  |  |  |
|  | Salivary glands | 27°C DTR 0°C | ZT12 | 120 | 14-16 | 4 | 150 or 120^ǂ^ |  |  |  |  |
|  |  |  | ZT18 | 120 | 14-16 | 4 | 150 or 120^ǂ^ |  |  |  |  |
|  |  |  | ZT0 | 120 | 14-16 | 4 | 150 or 120^ǂ^ |  |  |  |  |
|  |  | 27°C DTR 10°C | ZT12 | 120 | 14-16 | 4 | 150 or 120^ǂ^ |  |  |  |  |
|  |  |  | ZT18 | 120 | 14-16 | 4 | 150 or 120^ǂ^ |  |  |  |  |
|  |  |  | ZT0 | 120 | 14-16 | 4 | 150 or 120^ǂ^ |  |  |  |  |
| Supplementary Fig. 2a and Supplementary Table 4 (effects of time-of-day and fluctuating temperature on vector competence in *A. stephensi*) | Midguts | 27°C DTR 0°C | ZT12 | 60 | 7-9 | 2 | 120 | GLMM | Oocyst intensity or prevalence | Time-of-day + Temperature regime + Time-of-day × Temperature regime + Dissection day + Mosquito container^*^ | Oocyst intensity - negative binomial distribution with log link; Oocyst prevalence - binomial distribution with logit link |
|  |  |  | ZT18 | 60 | 7-9 | 2 | 120 |  |  |  |  |
|  |  |  | ZT0 | 60 | 7-9 | 2 | 120 |  |  |  |  |
|  |  | 27°C DTR 10°C | ZT12 | 60 | 7-9 | 2 | 120 |  |  |  |  |
|  |  |  | ZT18 | 60 | 7-9 | 2 | 120 |  |  |  |  |
|  |  |  | ZT0 | 60 | 7-9 | 2 | 120 |  |  |  |  |
|  | Salivary glands | 27°C DTR 0°C | ZT12 | 60 | 14-16 | 2 | 120 | GLM | Sporozoite prevalence^¶^ | Time-of-day + Temperature regime + Time-of-day × Temperature regime + Dissection day | Sporozoite prevalence - binomial distribution with logit link |
|  |  |  | ZT18 | 60 | 14-16 | 2 | 120 |  |  |  |  |
|  |  |  | ZT0 | 60 | 14-16 | 2 | 120 |  |  |  |  |
|  |  | 27°C DTR 10°C | ZT12 | 60 | 14-16 | 2 | 120 |  |  |  |  |
|  |  |  | ZT18 | 60 | 14-16 | 2 | 120 |  |  |  |  |
|  |  |  | ZT0 | 60 | 14-16 | 2 | 120 |  |  |  |  |
| Supplementary Fig. 2b and Supplementary Table 5 (effects of time-of-day and fluctuating temperature on vector competence in *A. stephensi*, a simplified version) | Midguts | 27°C DTR 0°C | ZT12 | 36 | 8-10 | 1 | 150 | GLM | Oocyst intensity or prevalence | Time-of-day + Temperature regime + Time-of-day × Temperature regime + Dissection day | Oocyst intensity - Poisson distribution^§^ with log link; Oocyst prevalence - binomial distribution with logit link |
|  |  |  | ZT23 | 31 | 8-10 | 1 | 150 |  |  |  |  |
|  |  | 27°C DTR 10°C | ZT12 | 30 | 8-10 | 1 | 150 |  |  |  |  |
|  |  |  | ZT23 | 32 | 8-10 | 1 | 150 |  |  |  |  |
|  | Salivary glands | 27°C DTR°0C | ZT12 | 31 | 13, 14, 16 | 1 | 150 | GLM | Sporozoite prevalence |  | Sporozoite prevalence - binomial distribution with logit link |
|  |  |  | ZT23 | 30 | 13, 14, 16 | 1 | 150 |  |  |  |  |
|  |  | 27°C DTR °10C | ZT12 | 28 | 13, 14, 16 | 1 | 150 |  |  |  |  |
|  |  |  | ZT23 | 30 | 13, 14, 16 | 1 | 150 |  |  |  |  |
| Fig. 3a and Supplementary Table 7 (effects of high temperatures on parasite establishment) | Midguts | 27°C | A. gambiae | 40 | 7-9 | 1 | 120 | GLM | Oocyst intensity or prevalence^¶^ | Oocyst intensity - Species + Temperature treatment + Species × Temperature treatment;  Oocyst prevalence - Temperature | Oocyst intensity - negative binomial distribution with log link; Oocyst prevalence - binomial distribution with logit link |
|  |  |  | A. stephensi | 30 | 7-9 | 1 | 120 |  |  |  |  |
|  |  | 30°C | A. gambiae | 25 | 6, 7 | 1 | 120 |  |  |  |  |
|  |  |  | A. stephensi | 28 | 6, 7 | 1 | 120 |  |  |  |  |
|  |  | 32°C | A. gambiae | 29 | 5, 6 | 1 | 120 |  |  |  |  |
|  |  |  | A. stephensi | 25 | 5, 6 | 1 | 120 |  |  |  |  |
| Fig. 3b (thermal sensitivity of early parasite infection in *A. stephensi*) | Midguts | 27°C | Control | 37 | 7-9 | 1 | 120 | GLM | Oocyst intensity or prevalence | Temperature treatment | Oocyst intensity - Poisson distribution^§^ with log link; Oocyst prevalence - binomial distribution with logit link |
|  |  | 30°C | 3h | 44 | 5-8 | 1 | 120 |  |  |  |  |
|  |  |  | 6h | 44 | 5-8 | 1 | 120 |  |  |  |  |
|  |  |  | 12h | 36 | 6-8 | 1 | 120 |  |  |  |  |
|  |  |  | 24h | 32 | 6-8 | 1 | 120 |  |  |  |  |
|  |  |  | 48h | 30 | 6-8 | 1 | 120 |  |  |  |  |
| Fig.4 (infectious feed for thermal avoidance assay) | Midguts | 27°C DTR 10°C prior to blood feeding, and 27°C after blood feeding at 06:00h (ZT0) | | 60 | 8 | 3 | 100 | NA | | | |
| Supplementary Fig. 3a and Supplementary Table 8 (effects of gametocytemia dilutions and high temperature on parasite establishment in *A. stephensi*) | Midguts | 27°C | 1 | 32 | 8, 9 | 1 | 120 | GLM | Oocyst intensity or prevalence | Gametocytemia + Temperature treatment + Gametocytemia × Temperature treatment | Oocyst intensity - negative binomial distribution with log link; Oocyst prevalence - binomial distribution with logit link |
|  |  |  | 1/2 | 33 | 8, 9 | 1 | 120 |  |  |  |  |
|  |  |  | 1/4 | 31 | 8, 9 | 1 | 120 |  |  |  |  |
|  |  |  | 1/10 | 31 | 8, 9 | 1 | 120 |  |  |  |  |
|  |  | 30°C | 1 | 48 | 7 | 1 | 120 |  |  |  |  |
|  |  |  | 1/2 | 55 | 7 | 1 | 120 |  |  |  |  |
|  |  |  | 1/4 | 54 | 7 | 1 | 120 |  |  |  |  |
|  |  |  | 1/10 | 50 | 7 | 1 | 120 |  |  |  |  |
| Supplementary Fig. 4 (effect of transferring mosquitoes between different temperatures on vector competence in *A. gambiae*) | Midguts | 27°C | | 30 | 8,9 | 1 | 120 | GLM | Oocyst intensity, or oocyst or sporozoite prevalence | Mosquito container | Oocyst intensity - negative binomial distribution with log link; Oocyst and sporozoite prevalence - binomial distribution with logit link |
|  |  | 21°C | | 60 | 14-16 | 2 | 120 |  |  |  |  |
|  | Salivary glands | 27°C | | 30 | 14-16 | 1 | 120 |  |  |  |  |
|  |  | 21°C | | 60 | 34-36 | 2 | 120 |  |  |  |  |
| Supplementary Table 9 and 10 (effect of blood feeding at different temperature on feeding compliance in *A. gambiae*) | NA | 27°C DTR 10°C | ZT12 | NA | NA | 2 | 120 | GLMM | Blood feeding success of individual mosquitoes | Time-of-day + Temperature regime + Time-of-day × Temperature regime + Mosquito container^*^ | Blood feeding success - binomial distribution with logit link |
|  |  |  | ZT18 |  |  | 2 | 120 |  |  |  |  |
|  |  |  | ZT0 |  |  | 2 | 120 |  |  |  |  |
|  |  | 21°C DTR 10°C | ZT12 |  |  | 2 | 120 |  |  |  |  |
|  |  |  | ZT18 |  |  | 2 | 120 |  |  |  |  |
|  |  |  | ZT0 |  |  | 2 | 120 |  |  |  |  |
| Supplementary Fig. 5 and Supplementary Table 11 (effect of transferring mosquitoes between different temperatures on blood meal size in *A. gambiae*) | NA | 27°C DTR 10°C | ZT12 | 20 | NA | 2 | 30 | GLMM | Mosquito body weight | Species + Time-of-day + Species × Time-of-day + Mosquito container^*^ | Mosquito body weight - normal distribution with identity link |
|  |  |  | ZT18 | 20 |  | 2 | 30 |  |  |  |  |
|  |  |  | ZT0 | 20 |  | 2 | 30 |  |  |  |  |

^†^Dpi: Days post infection

^ǂ^150 or 120 mosquitoes per container for each of two biological replicate experiments

^*^Included as a random variable in model analysis

^¶^Prevalence data were pooled within each temperature treatment group after confirming no difference between two replicates or species (Fisher’s exact test, two-sided, *P* > 0.05) to ensure model validity^44^^,^^45^

^§^Poisson distribution was used to ensure best model fit based on AIC value
